# Plant peptidoglycan precursor biosynthesis: Conservation between moss chloroplasts and Gram negative bacteria

**DOI:** 10.1101/2022.01.05.475093

**Authors:** Amanda J. Dowson, Adrian J. Lloyd, Andrew C. Cuming, David I. Roper, Lorenzo Frigerio, Christopher G. Dowson

## Abstract

An accumulation of evidence suggests that peptidoglycan, consistent with a bacterial cell wall, is synthesised around the chloroplasts of many photosynthetic eukaryotes, from glaucophyte algae to land plants at least as evolved as pteridophyte ferns, but the biosynthetic pathway has not been demonstrated. We employed mass spectrometry and enzymology in a twofold approach to characterize the synthesis of peptidoglycan in chloroplasts of the moss *Physcomitrium (Physcomitrella) patens*. To drive the accumulation of peptidoglycan pathway intermediates, *P.patens* was cultured with the antibiotics phosphomycin, D-cycloserine and carbenicillin, which inhibit key peptidoglycan pathway proteins in bacteria. Mass spectrometry of the TCA-extracted moss metabolome revealed elevated levels of five of the predicted intermediates from UDP-Glc*N*Ac through to the UDP-Mur*N*Ac-D,L-diaminopimelate (DAP)-pentapeptide.

Most Gram negative bacteria, including cyanobacteria, incorporate *meso*-diaminopimelate (D,L-DAP) into the third residue of the stem peptide of peptidoglycan, as opposed to L-lysine, typical of most Gram positive bacteria. To establish the specificity of D,L-DAP incorporation into the *P.patens* precursors, we analysed the recombinant protein, UDP-Mur*N*Ac-tripeptide ligase (*MurE*), from both *P.patens* and the cyanobacterium *Anabaena* sp. strain PCC 7120. Both ligases incorporated D,L-DAP in almost complete preference to L-Lys, consistent with the mass spectrophotometric data, with catalytic efficiencies similar to previously documented Gram negative bacterial MurE ligases. We discuss how these data accord with the conservation of active site residues common to DL-DAP-incorporating bacterial MurE ligases and of the probability of a horizontal gene transfer event within the plant peptidoglycan pathway.

## Introduction

The endosymbiotic theory for the origin of photosynthetic eukaryotes proposes that an engulfed cyanobacterium evolved into the first ancestors of chloroplasts (Dagan et al., 2013; Ponce-Toledo et al., 2017). As with bacteria, these organelles (cyanelles) were surrounded by a peptidoglycan (or murein) wall (Scott et al., 1984). In bacteria, peptidoglycan covers the organism in a mesh-like ‘sacculus’ confering resistance to osmotic stress, and a species-specific shape and size. Although originally considered likely that peptidoglycan was lost from all photosynthetic organelles immediately after the glaucophyte branch (Pfanzagl et al., 1996), there has been an accumulation of evidence including sensitivity of chloroplast division to peptidoglycan-directed antibiotics, fluorescent labelling studies and gene knockout phenotypes to indicate that many streptophytes, including the charophyte algae (Matsumoto et al., 2012; Takano et al., 2018) and some bryophytes and pteridophytes (sister lineages to seed plants) (Takano and Takechi, 2010; Hirano et al., 2016), may have chloroplasts that synthesize peptidoglycan. Furthermore, in gymnosperms (Lin et al., 2017) and also a diverse number of eudicots (van Baren et al., 2016) all the critical genes for peptidoglycan synthesis have been identified, although a potential penicillin binding protein (PBP) typically required for peptidoglycan cross-linking has not been confirmed in eudicots.

The earliest evidence for peptidoglycan in embryophytes was uncovered when antibiotics affecting bacterial peptidoglycan synthesis in the bryophyte moss *P. patens* (Kasten and Reski, 1997; Katayama et al., 2003) and lycophytes and ferns (Izumi et al., 2008) were found to cause a decrease in chloroplast number with the formation of giant (macro)chloroplasts. Subsequently, genomics and *in silico* analyses confirmed the presence of all essential bacterial genes for peptidoglycan biosynthesis (Rensing et al., 2008). These genes are nuclear-encoded, predominantly plastid-targeted (Machida et al., 2006; Homi et al., 2009) and transcribed, as revealed by expressed sequence tags (ESTs). More recently, a peptidoglycan layer surrounding *P. patens* chloroplasts has been visualized using a fluorescently labelled substrate (Hirano et al., 2016) and electron microscopy (Sato et al., 2017).

Peptidoglycan in Gram negative bacteria has a repeating disaccharide backbone of β-(1,4) linked N-acetylglucosamine (Glc*N*Ac) and N-acetylmuramic acid (Mur*N*Ac) to which is appended a stem peptide comprising L-Ala, D-Glu, D,L-DAP, D-Ala--D-Ala. Variations in the amino acid residues have been identified and are consequent on either the specificity of the Mur ligases (MurC-F) or later modifications in peptidoglycan biosynthesis. In Gram positive bacteria MurE typically incorporates L-Lys as opposed to D,L-DAP, although Bacilli are a notable exception and several other amino acids have been identified in this position (Schleifer and Kandler, 1972; Barreteau et al., 2008; Vollmer et al., 2008). The stem peptides of adjacent saccharide strands are crosslinked by transpeptidation to stabilize the mature peptidoglycan (see biosynthetic pathway Figure 1).

**Figure 1.**
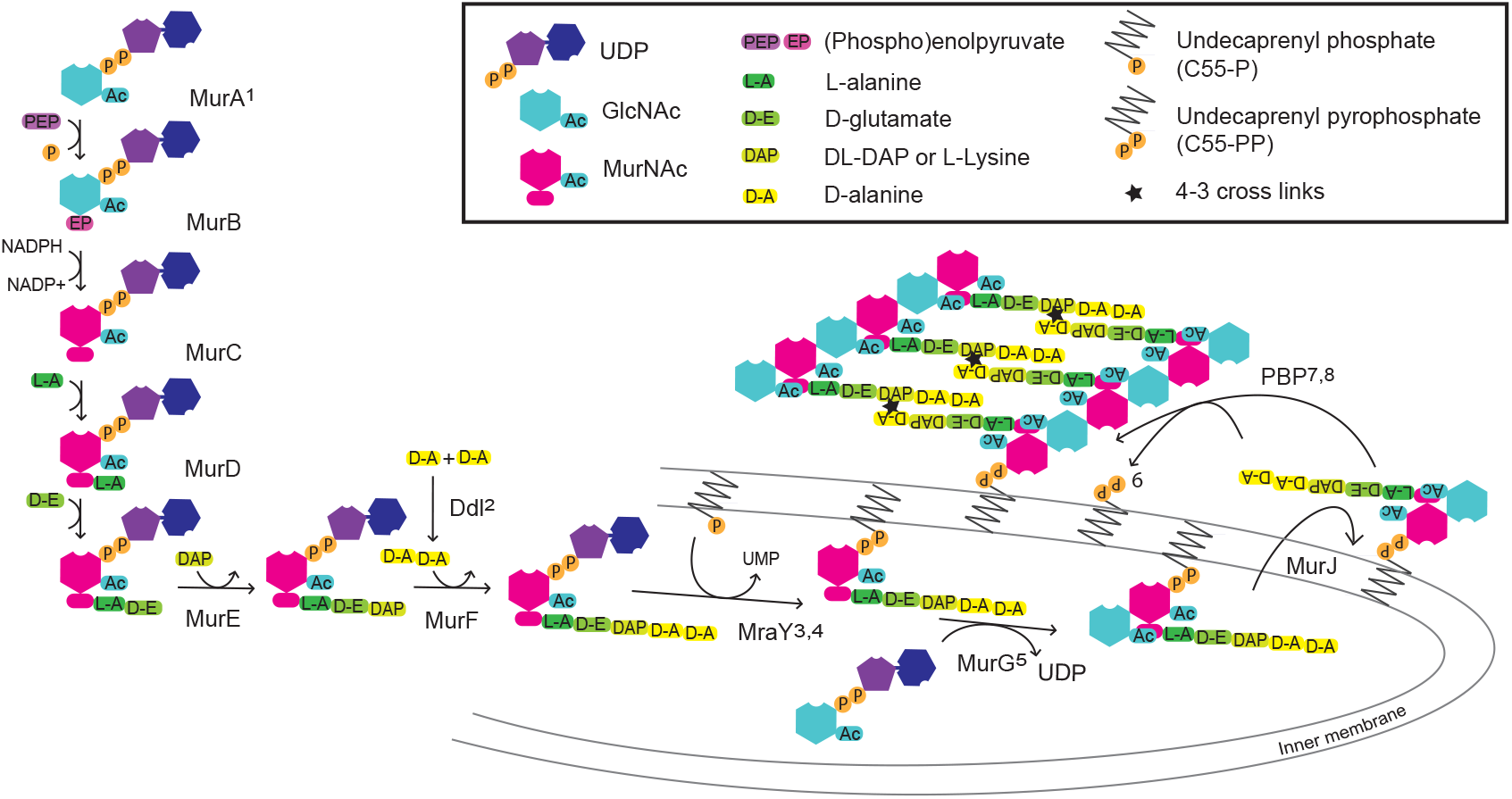
Schematic of the fundamental cytoplasmic and periplasmic enzyme steps in peptidoglycan (murein) biosynthesis. Enzymes: MurA-J, murein synthases A-J; Ddl, D-Ala--D-Ala ligase; MraY, phospho-N-acetylmuramoyl-pentapeptide-transferase and PBP, transglycosylase and transpeptidase activities of penicillin-binding proteins. Superscript numbers indicate targets for the following antibiotics: 1, phosphomycin, 2, D-cycloserine, 3, pacidamycin, 4, tunicamycin, 5, murgocil, 6, bacitracin, 7, penicillins and 8, vancomycin. The cytoplasmic Mur proteins MurA and MurB catalyze the formation of UDP-*N*-acetylmuramic acid (UDP-Mur*N*Ac), Mur ligases (MurC, D, E and F) sequentially append amino acids to form UDP-Mur*N*Ac-pentapeptide. The transmembrane protein MraY attaches Mur*N*Ac-pentapeptide to C55-P to yield C55-PP-Mur*N*Ac-pentapeptide (lipid I) and MurG Glc*N*Ac transferase creates C55-PP-Mur*N*Ac-(pentapeptide)-Glc*N*Ac (lipid II). Finally, the disaccharide pentapeptide monomer is flipped into the periplasm, polymerized by the transglycosylase activities of penicillin-binding-proteins (PBPs), or functionally related shape, elongation, division and sporulation (SEDS) proteins, and the peptides are 4-3 cross-linked to pre-existing peptidoglycan by the transpeptidase activities of PBPs or 3-3 cross-linked by L,D-transpeptidases. C55-PP is then subject to pyrophosphatase activity and C55-P recycled.

Knock-out of *P. patens* homologs of bacterial peptidoglycan synthesis genes *Ddl, MurA, MurE, MraY, MurJ* or *PBP1A,* results in a macrochloroplast phenotype, similar in appearance to antibiotic treatments that target their gene products, while complementation with the intact genes restores the wild type number of about 50 typical chloroplasts per cell (Machida et al., 2006; Homi et al., 2009; Hirano et al., 2016; Takahashi et al., 2016; Utsunomiya et al., 2020). Cross-species complementation using a *P.patens MurE* (*PpMurE)* knockout showed that *Anabaena MurE* (*AnMurE*) fused to the plastid-targeting signal of *PpMurE* can also restore the wild type chloroplast phenotype (Garcia et al., 2008). In contrast, the homologous *Arabidopsis thaliana gene*, *AtMurE*, failed to complement the *PpMurE* mutant (Garcia et al., 2008). Interestingly, *MurE* knockouts in both *A. thaliana* and *Zea mays,* appear bleached as opposed to having a macrochloplast phenotype, are deficient in chloroplast thylakoids and lack many plastid RNA polymerase-regulated chloroplast transcripts, indicating that angiosperm MurE has a primary function in plastid gene expression and biogenesis rather than plastid division *per se* (Garcia et al., 2008; Williams-Carrier et al., 2014).

Although data suggestive of the formation of chloroplast peptidoglycan is available, no direct observation of the peptidoglycan precursors or the operation of the chloroplast peptidoglycan synthetic pathway has yet been made. Therefore, here, using pathway-inhibiting antibiotics to drive the accumulation of peptidoglycan intermediates, we establish that in a basal land plant, *P. patens*, the six *Mur* genes and *Ddl* actively synthesize all the main precursors of the peptide stem of peptidoglycan. Furthermore, we show that the pentapeptide building blocks are identical to those of most typical Gram negative bacteria, including the cyanobacteria, plus the Chlamydiae, the ‘acid fast’ *Mycobacterium* spp. and some Gram positive bacilli, where D,L-DAP is incorporated instead of L-Lys. Consistent with and supportive of this observation, we show that *in vitro* the moss MurE ligase, PpMurE, incorporates D,L-DAP in strict preference to L-Lys as the third amino acid within the stem peptide, as would be consistent with the cyanobacterial ancestral origin of the chloroplast, and the enzyme kinetics of PpMurE are similar to cyanobacterial and other Gram negative D,L-DAP-incorporating MurE ligases.

## Materials and Methods

### Plant Material

*P. patens* (Gransden strain, GrD13) was grown on modified KNOPS medium with 5 mM diammonium tartrate, to promote chloronemata formation (Schween et al., 2003). The medium was solidified with 0.85 % (w/v) plant agar (Sigma) and overlaid with 9 cm cellophane discs (AA Packaging). Plants were grown in 90 mm diameter x 20 mm vented tissue culture dishes sealed with Micropore (3M) surgical tape in a plant growth room at 21°C under continuous light from Sylvania white F100W tubes at 65-100 μmol.m^−2^.s^−1^. After being homogenised axenically in water in a 250 ml flask using an IKA T18 digital Ultra Turrax homogeniser, for one to two 12 s bursts, *P. patens* protonemata were cultured as 2 ml aliquots per 25 ml solid KNOPS plus tartrate.

### Confocal Microscopy of Antibiotic Treated *P. patens* Protonemata

Confocal single plane images and Z-series stacks were acquired on a Leica SP5 microscope, using a 63 x 1.4 Oil UV immersion objective with the 405 nm and 496 nm laser lines and transmitted light, and photo multiplier tube spectral detection adjusted for the chlorophyll emission (735-790 nm). Images were processed using the Fiji distribution of ImageJ v2.0.0.

### Trichloroacetic Acid (TCA) Extraction of Plant Metabolites

Antibiotics were added to KNOPS plus tartrate agar at 100 µg ml^-1^ carbenicillin, 100 µg ml^-1^ D-cycloserine or 200 µg ml^-1^ phosphomycin. After 15 days tissue was harvested, weighed and ground in liquid nitrogen using a pestle and mortar before being frozen at −80°C. To extract TCA-soluble plant metabolites the tissue was ground again in 5 ml g^-1^ of ice cold 10% (w/v) TCA (Fisons AR grade) before being mixed gently in 50 ml Falcon tubes on a rolling shaker for 30 min at 4°C (Roten et al., 1991). Insoluble material was pelleted at 48,000 xg, 10 min, 2°C, the supernatant was retained and the pellet reextracted twice more, first with 2.5 ml.g^-1^ and then with 1.25 ml.g^-1^ (of the original pellet weight) of ice cold 10% (w/v) TCA. The pooled supernatants were extracted into an equal volume of diethyl ether, to remove TCA, by manually shaking for 3 x 20 s in a separating funnel before recovering the lower, aqueous layer. The ether extraction of the aqueous phase was repeated twice more. The pH of the combined lower phases was restored to pH 7-8 using 1 M NaOH and residual ether was removed *in vacuo* at which point, the sample was lyophilised.

### Purification of Muropeptide Precursors

The nucleotide precursors in the TCA-soluble metabolite extracts were first partially purified by size exclusion using a Superdex Peptide 10/300GL column. The freeze-dried pellets were resuspended in deionised water, applied to the column as a 500 µl aliquot, eluted with deionised water and collected as 0.5 ml fractions at 0.5 ml min^-1^. The likely elution volume of molecules of interest was established by elution of 20 nmols UDP-Mur*N*Ac-DAP-pentapeptide and 20 nmols UDP-*N*-acetyl-glucosamine (Sigma) standards.

The *A*_260_ of pooled Superdex Peptide fractions of 1-2 ml was used to determine the upper linit of the total concentration of UDP species and an estimated 2, 10 or 20 nmols UDP species in 2 ml 10 mM ammonium acetate, pH 7.5, was loaded onto a MonoQ 50/5 GL column equilibrated in the same buffer. Bound molecules were eluted with a 27 ml linear gradient of 10 mM to 0.81M ammonium acetate (pH 7.5), at 0.7 ml.min^-1^ and collected as 1ml fractions using an Äkta Pure where the eluate absorbance was recorded at *A*_230_, *A*_254_ and *A*_280_. Peaks with an absorbance ratio of usually 1:2 A_280_:A_254_ were selected for freeze drying and mass spectrometry.

### Mass Spectrometric Nano-spray Time of Flight Analysis of Peptidoglycan UDP-MurNAc Precursors

Identity of UDP-MurNAc precursors were confirmed by negative ion time of flight mass spectrometry using a Waters Synapt G2Si quadrupole-time of flight instrument operating in resolution mode, equipped with a nanospray source calibrated with an error of less than 1 ppm with sodium iodide over a 200-2500 m/z range (Catherwood et al., 2020). Samples, freeze dried three times to remove ammonium acetate, were diluted in LCMS grade 50% v/v acetonitrile to between 1 µM and 5 µM. They were introduced into the instrument using Waters thin wall nanoflow capillaries and up to 20 minutes of continuum data were collected at a capillary voltage of 2.0 kV, cone and source offset voltages of 100 V and a source offset of 41 V, respectively. Source and desolvation temperatures were 80°C and 150°C respectively, desolvation and purge gas flow rates were both 400 l.min^-1^. Scan time was 1 s with an interscan time of 0.014 s. Scans were combined into centred mass spectra by Waters Mass Lynx software. Resolution (m/z/half-height spectral peak width) was measured as 1 in 20,100.

### Construction of Heterologous Expression Plasmids

*PpMurE* (derived from Pp3c23_15810V3.2) and *AnMurE* (derived from *Anabaena* sp. strain PCC 7120 *MurE* WP_010995832.1 Q8YWF0|MURE_NOSS1) were inserted into the vector pPROEX HTa (Addgene) in order to be expressed in frame with an amino terminal, TEV protease-cleavable, hexa-histidine (His6) tag. The *MurE* sequences were PCR amplified from their respective cDNAs (Machida et al., 2006; Garcia et al., 2008) in pTFH22.4 using the primers PpMurE_L63_Forward (TTTGCGACATGTTGAAAATGGGGTTTGGGGATTCGAAATTGACGGATCG) and PpMurE_Reverse (AAACGCGCGGCCGCTTATTTTCTAAGTCGCAAAGCCTCCCGACATTCCTC) and Anabaena_PCC7120_MurE_Forward (TTTGCGGGTCTCTCATGAAATTGCGGGAATTACTAGCGACAGTAGACAGTG) and Anabaena_PCC7120_MurE_Reverse (AAACGCGCGGCCGCTTATAATTTTTCTCTTTCTGTCAAAGCGGCGCGTGCG). The amplified region for *PpMurE* started at leucine 63, effectively deleting the chloroplast transit peptide at the cleavage site predicted by the ChloroP1.1 Prediction server (Emanuelsson, 1999) and introducing a unique *Nco1*-compatible *Pci1* site around the novel ATG and a *Not1* site immediately 3’ to the stop codon. A TAA stop codon was substituted for the native TGA. The *AnMurE* primers amplified the cDNA and novel *Bsa1* and *Not1* sites were introduced 5’and 3’ to the ATG start and TAA stop codons, respectively. The former was sited to create a *Nco1*-compatible 5’ cohesive end. The vector pPROEX HTa was restricted with *Nco1* and *Not1* and gel purified before being ligated to *Pci1-Not1* restricted *PpMurE_L63* or *Bsa1-Not1* restricted *AnMurE* PCR fragments that had been cleaned up with a PCR clean up kit (Qiagen). Coding sequences were confirmed by Sanger sequencing (Eurofins).

### Expression of *PpMurE_L63* and *AnMurE* and Protein Purification

For protein purification *Escherichia coli* strains were tested for optimal expression: BL21 DE3 (Thermofisher) was selected for PpMurE_L63_pPROEX and BL21(DE3), with the chaperone plasmid pG-KJE8 (Takara Bio Inc.), was selected for AnMurE_pPROEX. These were grown in L-Broth plus 0.2% v/v glucose, 100 µg ml^-1^ ampicillin and 35 µg ml^-1^ chloramphenicol at 37°C to an *A*_600_ of 0.6 when PpMurE protein expression was induced with 0.5 mM IPTG and AnMurE expression was induced by 0.5 mM IPTG with 1.5 mg.ml^-1^ arabinose and 8 ng.ml^-1^ tetracycline to induce pG-KJE8 chaperones. Over-expressing cells were then grown overnight at 19°C. Bacteria were harvested by centrifugation at 5600 x g, 15 min at 4°C and resuspended in Buffer A: 50 mM HEPES-NaOH, 0.5 M NaCl, 10 mM imidazole and 10% v/v glycerol (pH 7.5) containing EDTA-free protease inhibitor tablets, as recommended by the supplier (Pierce), and 2.5 mg ml^-1^ lysozyme, with gentle mixing for 30 min at 4°C. Lysis was by sonication on ice for 10 x 15 s bursts at 70%, interspersed by 1-2 min cooling on ice. Insoluble material was pelleted at 50,000 xg for 30 min at 4°C and the supernatant loaded directly onto a 5 ml His Trap HP (GE Healthcare) at 2 ml.min^-1^ and washed with 50 ml Buffer A at 4 m.min^-1^ at 4°C. Bound material was eluted with an 100 ml linear gradient to 100% Buffer B: 50 mM HEPES-NaOH, 0.5 M NaCl, 5% w/w glycerol and 0.5 M imidazole (pH 7.5) at 4 ml.min^-1^. Selected peak fractions were pooled and concentrated in either 30 or 50 kDa MWCO Vivaspin concentrators (GE Healthcare), for AnMurE or PpMurE_L63, respectively, at 2,800 xg at 4°C. Proteins were further purified by size exclusion chromatography on Superdex G200 XK26 (GE Life Sciences) pre-equilibrated and eluted with 50 mM HEPES-NaOH, 150 mM NaCl (pH 7.5) and purity of the eluted MurE proteins was established by SDS-PAGE (Supplemental Figure S3). Pooled peak fractions were dialysed against DB2: 30 mM HEPES-NaOH, 1 mM MgCl_2_, 50 mM NaCl, 50% v/v glycerol with 0.2 mM PMSF, 1 µM leupeptin, 1 µM pepstatin, 3 mM dithiothreitol (pH 7.6) overnight at 4°C, before storage at −20°C and −80°C.

### TEV Protease-Cleaved Protein Preparation

Bacteria were expressed as above, harvested and lysed using a cell disruptor and the proteins first purified on 5ml His Trap HP columns, using Buffer A and B (pH 8.0) as above, except that 100 mM Tris replaced 50 mM HEPES and Buffer A included 2% v/v glycerol, 10 mg.l^-1^ DNase1 (DN25) and 1 mM DTT. Pooled fractions were exchanged into a buffer of 50 mM PIPES, 100 mM NH_4_SO_4_, 200 mM KCl, 20 mM MgCl_2_, 1 mM DTT, 30 mM imidazole, 2% v/v glycerol (pH 7.7). using a stack of four 5ml HiTrap Desalting columns (Pharmacia). Peak fractions were incubated for 48h at 4°C in the ratio 1 mg TEV protease: 50 mg protein before reverse His-tag purification, collecting the column flow through. Samples were concentrated using 50 kDa concentrators as above.

*Streptococcus pneumoniae* MurE and *Pseudomonas aeruginosa* MurF were over-expressed and purified exactly as described (Blewett et al., 2004; Majce et al., 2013).

### Mur Ligase Activity Assays

The assays employed a continuous spectrophotometric method following ATP consumption at 37°C in a Cary 100 UV/Vis double beam spectrophotometer. Mur ligase catalysed ADP release, coupled to NADH oxidation by pyruvate kinase and lactate dehydrogenase, led to stoichiometric consumption of NADH measured by a fall in the *A*_340_. Assay volumes were 0.2 ml and contained 50 mM PIPES, 10 mM MgCl_2_ adjusted to pH 6.7 for AnMurE or 50 mM Tricine, 10 mM MgCl_2_ adjusted to pH 8.7 for PpMurE_L63, 1 mM dithiothreitol, 0.2 mM NADH, 2 mM phospho*enol* pyruvate, 1mM ATP, 50 mM.min^-1^ pyruvate kinase and 50 mM.min^-1^ lactate dehydrogenase (as assayed by the manufacturer, Sigma). Ligases were diluted prior to assay as required in 50 mM HEPES pH 7.7, 50 mM KCl, 1 mM MgCl_2,_ 3 mM DTT, 50% v/v glycerol, 0.2 mM PMSF. Concentrations of UDP-Mur*N*Ac dipeptide, Mur ligase and amino acid substrates were as described in the text or table legends. Control rates were collected usually in the absence of the amino acid, or UDP-Mur*N*Ac-dipeptide as specified, and the activity of the enzyme was initiated by addition of the missing component. Mur ligase initial rates were recorded as mols ADP.mol Mur ligase^-1^.s^-1^ (ADP/s) within the linear range of the time course of the assay.

## Results

### Identification of antibiotics with the most profound effect on *P. patens* chloroplast division

*P.patens* was grown on a variety of antibiotics that impact peptidoglycan biosynthesis in bacteria in order to select those best able to cause an accumulation of peptidoglycan intermediates in the moss, so that they might be more readily detected. Of the antibiotics tested the three that appeared most specific at inhibiting peptidoglycan synthesis, as measured by a widespread macrochloroplast phenotype with least effect on chlorophyll intensity, were phosphomycin (500 µg.ml^-1^), a PEP analog that inhibits MurA by alkylating an active site cysteine residue (Figure 1^1^), the ß-lactam ampicillin (100 µg.ml^-1^), which binds covalently to the active site serine of PBPs (Figure 1^7^), and D-cycloserine (20 µg.ml^-1^), with at least two enzyme targets in peptidoglycan biosynthesis, DDL (Figure 1^2^) and alanine racemase (Figure 2 B, D and G). Even at higher concentrations, where growth rate was impaired, the protonemata were green indicating chlorophyll synthesis and therefore chloroplast function was not significantly impaired. The impact of antibiotics that had either a more profound and pleiotropic effect or that had little impact on phenotype are described in Supplemental Text S1 and include vancomycin, bacitracin, murgocil and A22 (Figure 2 C, E, F and H).

**Figure 2.**
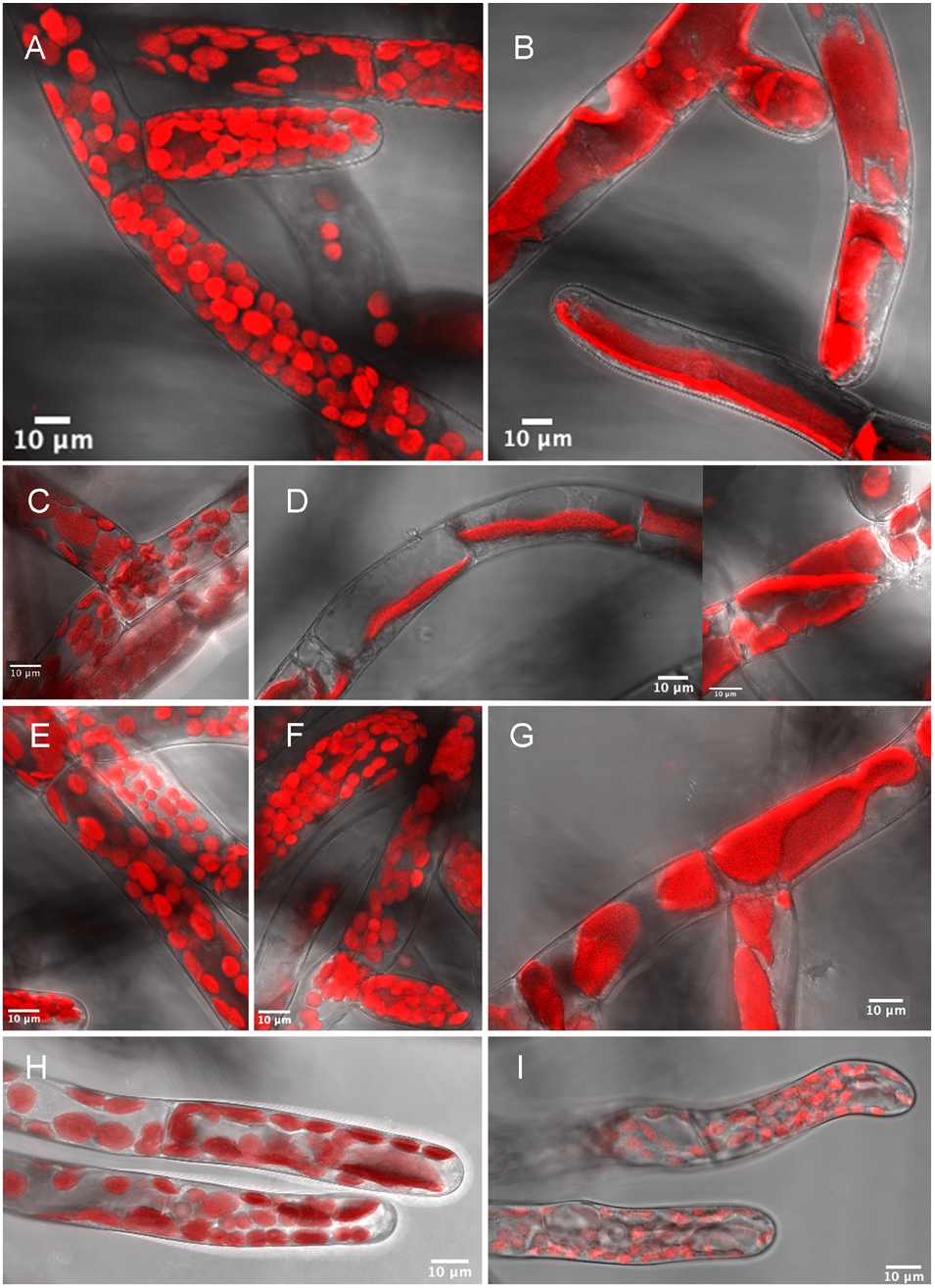
Confocal microscope images showing the effects of antibiotics on *P. patens* chloronemata. Chlorophyll autofluorescence (red) reveals macrochloroplasts consequent on growth on phosphomycin, D-cycloserine, vancomycin, bacitracin, ampicillin and A22. A. untreated, B. phosphomycin (500 µg.ml^-1^), C. vancomycin (25 µg.ml^-1^), D. D-cycloserine (20 µg.ml^-1,^ two images), E. bacitracin (100 µg.ml^-1^), F. murgocil (10 µg.ml^-1^), G. ampicillin (100 µg.ml^-1^), H. A22 (2.5 µg.ml^-1^) and I. A22 (10 µg.ml^-1^). Sequential fluorescence and transmitted light images, from a Leica SP5 with 63 x oil immersion lens, were processed using LAS AF lite to optimise intensity and combined as hyperstacks using Fiji on Image J. Scale bars 10µm.

### The TCA-extracted metabolome contains peptidoglycan precursors in *P. patens*

*P. patens* was grown separately on the three most specific and effective antibiotics, phosphomycin (400 µg.ml^-1^), D-cycloserine (100 µg.ml^-1^), and carbenicillin (100 µg.ml^-1^) to facilitate accumulation of different peptidoglycan precursor molecules (Figure 1^1, 2 and 7^). After size exclusion and anion exchange chromatography to purify UDP-linked intermediates from the TCA-extracted metabolome, mass spectrophotometric analysis identified precursors common to most Gram negative bacterial cell wall syntheses (Table 1 and Figure 3 C, identified precursors numbered 1-5). Precursor molecules were detected only in the earlier fractions from the Superdex Peptide column (Figure 3 B), as expected from the elution profiles of UDP-Glc*N*Ac and UDPMur*N*Ac-pentapeptide standards (not shown).

**Table 1.**
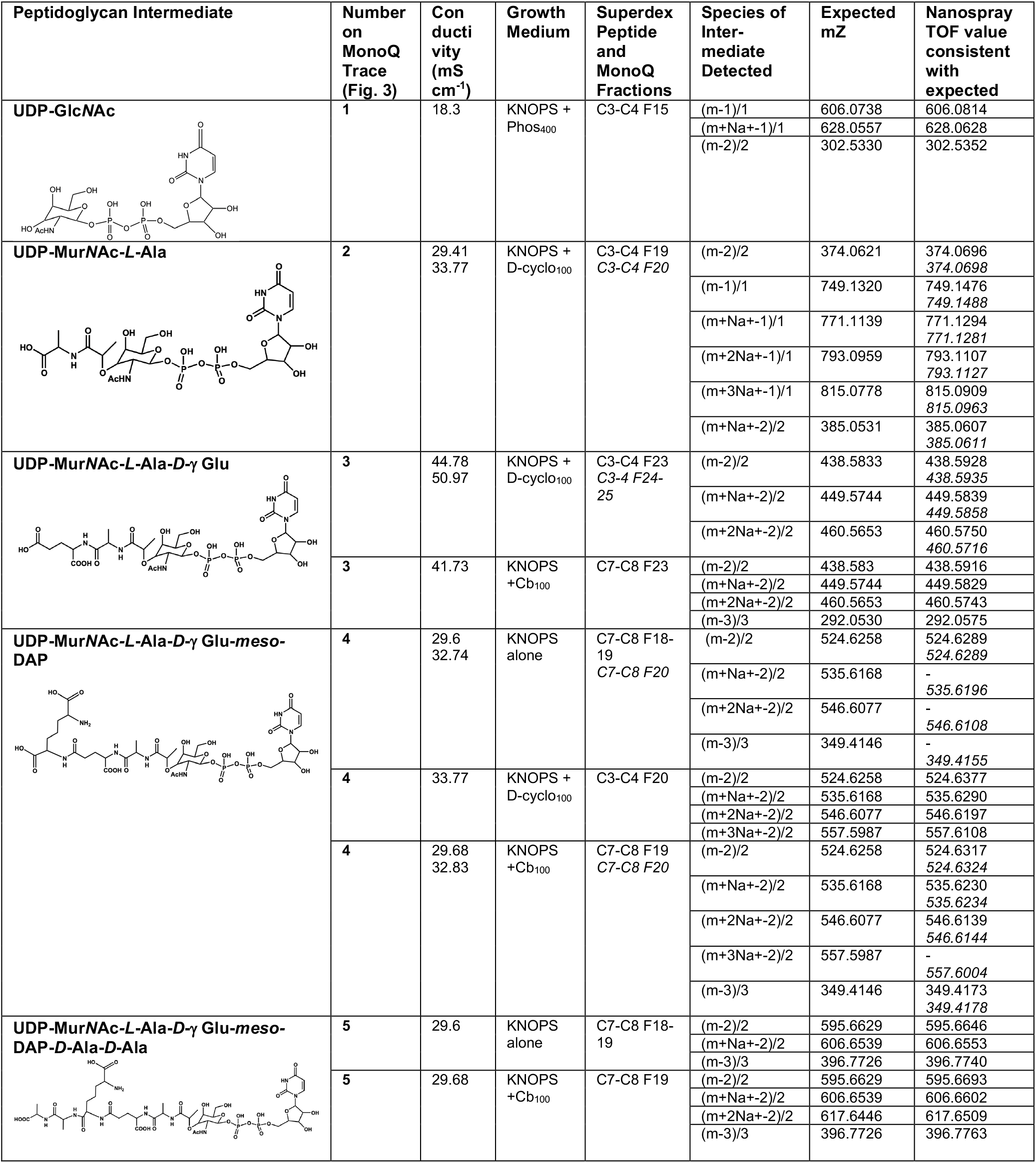
UDP-linked intermediates in peptidoglycan biosynthesis as detected by mass spectrometry of the *P. patens* TCA-extracted metabolome, with expected mass:charge (mZ) ratios and actual TOF nanospray values as listed. (Figures in italics represent where a species was detected in more than one fraction). *P. patens* was grown on KNOPS medium with or without antibiotics, including Phos_400_ (phosphomycin 400 μg.ml^-1^), D-cyclo_100_ (D-cycloserine 100 μg.ml^-1^) and Cb_100_ (carbenicillin 100 μg.ml^-1^). Superdex Peptide (C) and MonoQ fractions (F) where the different species were identified are listed with their peak conductivities on MonoQ, as detailed in Figure 3. The negative ion nanospray TOF mass spectra from which the data are derived are in Supplemental Figure S2.

**Figure 3.**
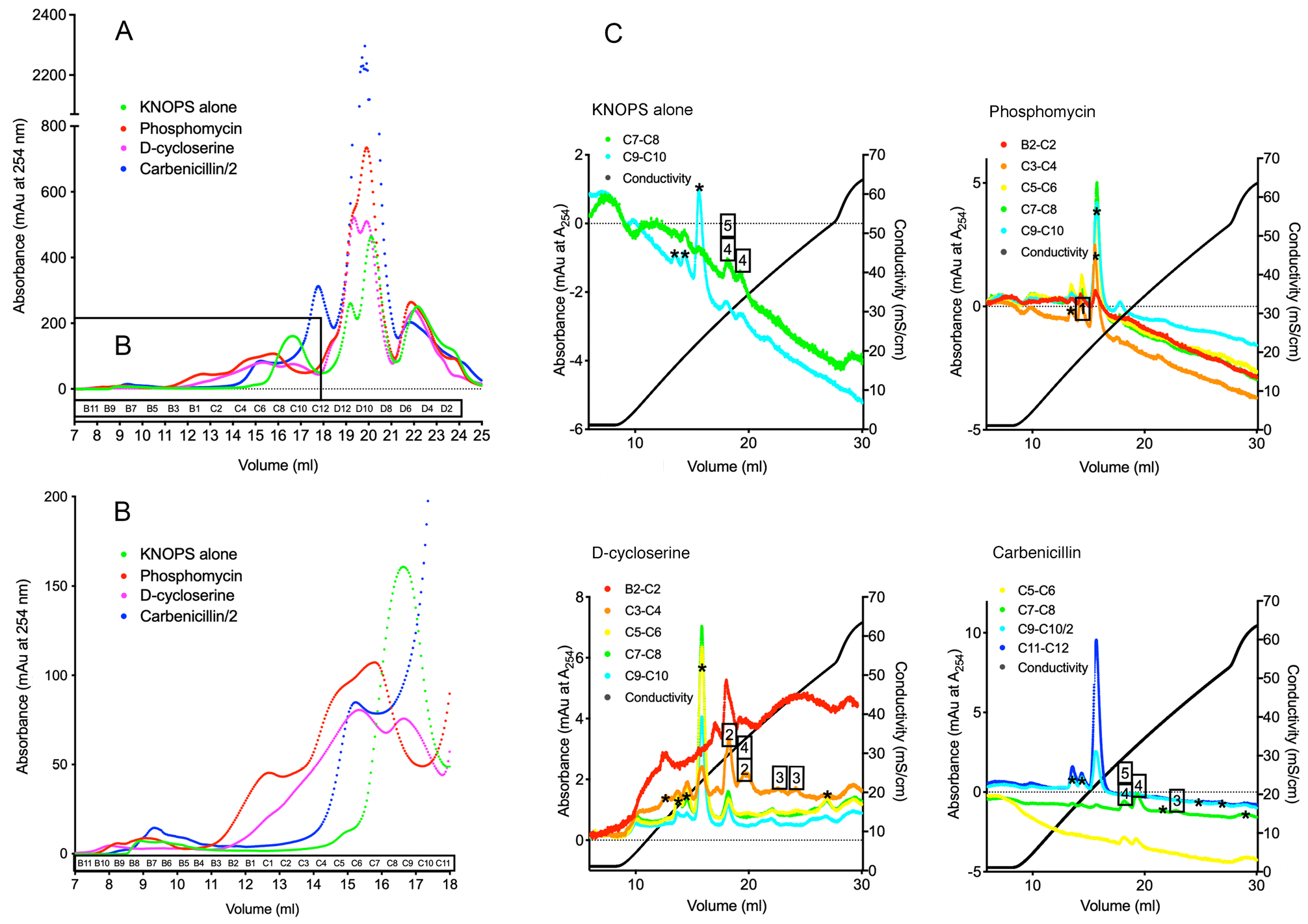
Size exclusion (Superdex Peptide, A and B) and ion exchange (MonoQ, C) chromatography elution profiles at A254 of the TCA-extracted metabolome from *P patens* grown on KNOPS with and without antibiotics. A. Superdex Peptide traces for the four treatments; KNOPS alone, or KNOPS with phosphomycin (400 µg.ml·^1^), D-cycloserine (100 µg.ml·^1^) or carbenicillin (100 µg.ml·^1^). (For carbenicillin the A254 was divided by two). B. Enlargement of the earlier fractions, where the intermediates were anticipated to elute (as determined by controls). C. MonoQ traces of pooled Superdex Peptide fractions of 2-20 nmoles of UDP species (from B2-C12). Boxed numbers represent fractions positively identified as intermediates (see Table 1) and asterisks indicate peaks with no recognised component.

The identification of UDP-Mur*N*Ac-Ala-Glu-D,L-DAP in three of the samples as well as the D,L-DAP pentapeptide (Table 1 and Figure 3 C, numbers 4 and 5), together with the inability to identify UDP-Mur*N*Ac-Ala-Glu-Lys or UDP-Mur*N*Ac-Lys-pentapeptide suggested that *in vivo*, PpMurE specifically incorporated DL-DAP in the third position of stem peptide. By comparison, when the plant was grown on phosphomycin (Figure 1^1^), anticipated to block synthesis of UDP-Mur*N*Ac, only the UDP-Glc*N*Ac precursor was identified (Table 1 and Figure 3 C, number 1). Interestingly, this metabolite was not detected in the samples treated with the other antibiotics.

Similarly, the MurC and D products, UDP-Mur*N*Ac-Ala and UDP-Mur*N*Ac-Ala-Glu were detected in the D-cycloserine-grown extract consistent with the accumulation of precursors up to the UDP-Mur*N*Ac-tripeptide MurF substrate (Figure 3 C, numbers 2 and 3). From the MonoQ anion exchange chromatograms (Figure 3, C) and the mass spectral data (Supplemental Figure S2) we can conclude that use of the different antibiotics proved to be an effective way to ensure most of the intermediates were detected, confirming the utility of this method for the purpose.

### *P. patens* MurE incorporates DL-DAP into the peptidoglycan stem peptide

To account for the composition of the *P.patens* peptidoglycan stem peptide, we analysed the activity and substrate specificity of the MurE ligase protein product of its *murE* gene, with the predicted 62 residue chloroplast transit peptide sequence deleted (PpMurE_L63). The enzyme was compared with the cyanobacterial *Anabaena* MurE ligase. Analysis of the ability of both AnMurE and PpMurE_L63 to utilise D,L-DAP D,D-DAP L,L-DAP and L-Lys revealed that both MurE enzymes were catalytically active in the aminoacylation of UDP-Mur*N*Ac-dipeptide, Removal of the His tag by TEV protease cleavage did not enhance the efficiency of either enzyme (Figure 4, B and Supplemental Figure S4) and significantly, both proteins favoured D,L-DAP as a substrate over the other DAP diastereoisomers (Figure 4, A). Noticeable was the slow rate of turnover of D,D-DAP by PpMurE_L63, in particular, possibly indicative of a weak stereo-selectivity for the L- over the D stereocentre of DAP utilised by the enzyme when at high concentrations. Significantly, neither enzyme incorporated L-Lys. As a control, lysylation of UDP-Mur*N*Ac-Ala-Glu was also assayed with the L-Lys specific *Streptococcus pneumoniae* Pn16 MurE (Blewett et al., 2004) and resulted in a rate (vo) of 1.94 ADP.s^-1^ at 150 µM L-Lys, with the same UDP-Mur*N*Ac-dipeptide and ATP concentrations as the other assays (data not shown).

**Figure 4.**
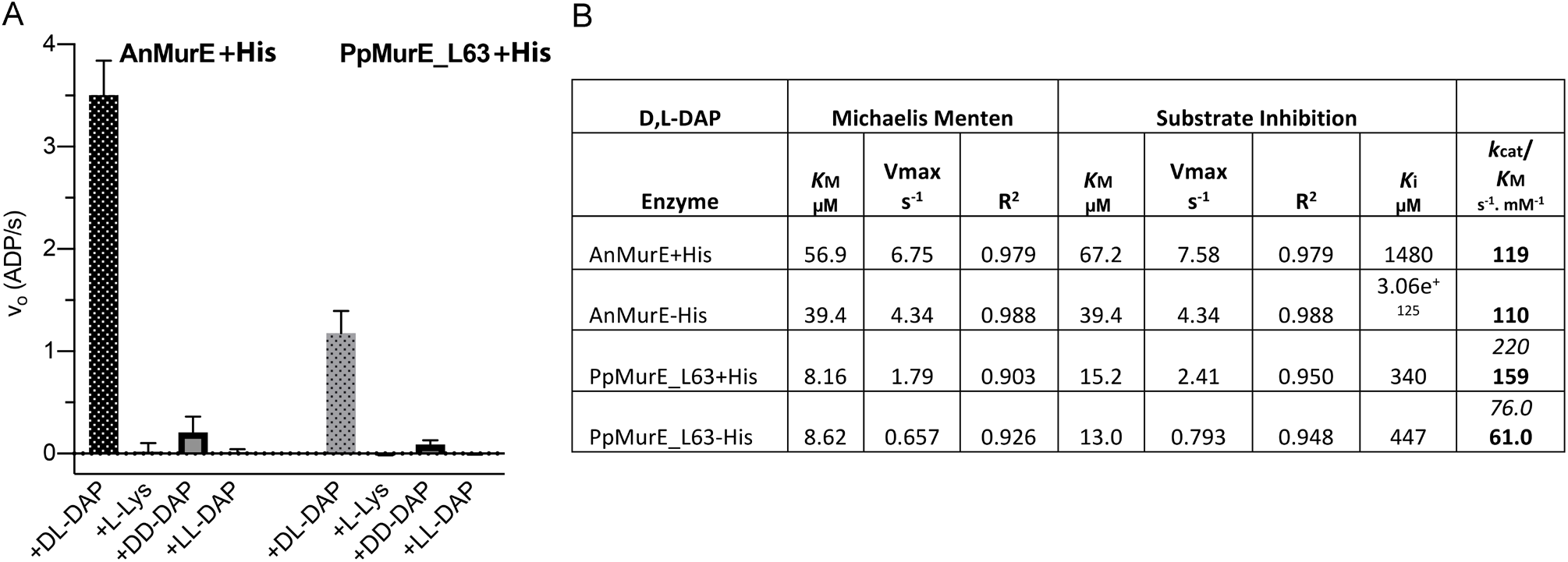
Substrate specificity and kinetics of AnMurE and PpMurE _L63. A, Activity of AnMurE and PpMurE_L63 with 150 µM D,L-DAP, L-Lys, D,D-DAP or L,L-DAP. Assays included 375 µM UDPMurNAc-dipeptide and 100 or 300 nM AnMurE+His or PpMurE_L63+His in 50 mM Hepes pH 7.6, respectively. Results (v) are presented as ADP.s·^1^ (molsADP.mol Mur ligase·^1^.s·^1^). Error bars are 95% confidence intervals of 5 or more rates from up to 8 replicate experiments. Students t test indicate D,D-DAP rates for both enzymes are significantly greater than those for either L,L-DAP or L-Lys. B. Michaelis Menten and substrate inhibition values for KM (µM), Vmax (ADP.s ^1^) and R^2^ (coefficient for data fit to either model), as computed by Prism, for both enzymes with and without His tags. All constants are ‘apparent’, obtained at fixed concentrations of the other two substrates. *K*cat derives from Vmax in mols ADP.mol Mur ligase·^1^.s·^1^. *K*cat*/K*M values are for Michaelis Menten kinetics for AnMurE and for substrate inhibition (bold) and Michaelis Menten kinetics (italics) for PpMurE_L63. (D,L-DAP substrate curves are in Supplemental Figure S4).

That the assay followed the aminoacylation of UDP-Mur*N*Ac-dipeptide by D,L-DAP to yield D,L-DAP tripeptide was confirmed by the ability of the assay product to be utilised as a substrate by *Pseudomonas aeruginosa* MurF (PaMurF). This was achieved in the same coupled assay by adding PaMurF at t=0, initiating the MurE ligase reaction with D,L-DAP and then the MurF ligase with D-Ala-D-Ala as the second substrate once the MurE reaction had reached completion to yield the UDP-MurNAc-pentapeptide (Supplemental Figure S5).

### pH and Temperature Optima of *P. patens* and *Anabaena* MurE

Prior to kinetic investigation of the properties of PpMurE the pH optimum was determined, with that of the cyanobacterial AnMurE, by comparing rate of ADP generated (vo) at pH 5.7-9.7 at approximately saturating concentrations of its substrates (Supplemental Figure S6, A-D). Neither of the coupled enzymes in the MurE/ADP release assay was a major factor affecting rate over the pH range studied as evidenced by the independence of the measured MurE rate from coupling enzyme concentration. Additionally, the similarity of activities of the MurE proteins in different buffers allowed us to discount the impact of buffers over the pH range under consideration (Supplemental Figure S6, B and D). Assuming saturation with substrates and the only variable responsible for a change in enzyme activity was pH range we fitted vo data vs pH to an equation that follows the relationship of activity *versus* pH. From these data it was concluded that the optimum for AnMurE is 7.5 and that for PpMurE_L63 is approximately pH 7.5-8.5. The data fit for for PpMurE_L63 (R^2^ = 0.94 and 0.89, for 1 and 2 x coupling enzymes, respectively) is better than that for AnMurE (R^2^ = 0.78) indicating that the assumption that other variables (kinetic constants and substrate ionization) are not influenced by pH may be less true for AnMurE.

### *P. patens* MurE has similar kinetic properties to cyanobacterial MurE

The two enzymes AnMurE and PpMurE_L63 were assayed to calculate their kinetic efficiency for the preferred substrate D,L-DAP. From the tabulated data PpMurE_L63 was more sensitive to substrate inhibition from D,L-DAP than AnMurE, as indicated by its greater R^2^ value for fit of data to the kinetics of substrate inhibition compared to those for standard Michaelis Menten kinetics (Figure 4, B and the two fitted curves in Supplemental Figure S4, C and D). However, the *K*cat^App^_/_*K*M^App^ ratio for the plant enzyme were similar to the cyanobacterial one, the most marked difference being the lower D,L-DAP *K*M^App^ value, indicative that the plant enzyme may operate at lower substrate concentrations *in vivo*. These figures were compared with reported data for other MurE activities (Supplemental Figure S7) and reveal that the plant and cyanobacterial MurE are at least as catalytically active, as indicated by the *K*cat^App^ *K*M^App^ ratio, as the bacterial homologs. It was apparent that removal of the His tag by TEV protease cleavage did not enhance the efficiency of either enzyme (Figure 4, B and Supplemental Figure S4, B and D).

### Conservation of amino acid residues common to DL-DAP-incorporating MurE ligases

BLASTP searches and ClustalW (EMBL-EBI) alignment indicated that the closest bacterial homolog to PpMurE is the MurE of *Gemmatomonidates bacterium* (50.0% homology), which is a photoheterotrophic Gram negative bacterium in a phylum quite distal to the cyanobacteria (Zeng et al., 2014). The next closest is the Gram positive *Bacillus fortis* (43.0%), which would be anticipated to incorporate D,L-DAP (Barreteau et al., 2008). Both are considerably more closely related than the cyanobacterial AnMurE (37.8%), determined in this paper to be D,L-DAP incorporating, *E.coli* MurE^D,L-DAP^ (34.9%) and *Mycobacterium tuberculosis* MurE^D,L-DAP^ (34.7%). The L-Lys incorporating enzymes, all from Gram negative species, share still less homology: *Thermatoga maritima* MurE^L-Lys^ (33.0%), *Streptococcus pneumoniae* MurE (30.1%) and *Staphylococcus aureus* MurE^L-Lys^ (26.6%). Likewise, the neighbour-joining phylogram computed in Jalview (Supplemental Figure S8) placed AnMurE as more distantly related than *Gemmatimonadetes* to plant MurE, as represented by PpMurE and the algal streptophytes *Mesotaenium endlicherianum* MurE and *Coleochaete scutata* MurE. *M. endlicherianum* (66.2%) represents a late charophyte ancestor within the Zygnemophyceae which are predicted to be on a branch point preceding embryophyte evolution (Donoghue and Paps, 2020), whereas MurE from *Klebsormidium nitens* (41.8%) and *C. scutata* (51.3%) in the Klebsomidiophyceae and Coleochaetaceae, respectively, and also within the charophyte algae, are on more divergent branches.

To relate homology to functionality, PpMurE was aligned in Clustal Omega (EMBL-EBI) with homologs of both L-Lys- and DL-DAP-incorporating MurE ligases (Supplemental Figure S9). Many amino acid residues are conserved not only between MurE from bacterial and early plant species but also across the Mur ligase family (as indicated by asterisks on Supplemental Figure S9). Mur ligases comprise three domains: an N-terminal Rossmann-fold domain responsible for binding the UDP-Mur*N*Ac substrate; a central ATP-binding domain and a C-terminal domain associated with binding the incoming amino acid. Most of the amino acids conserved between the different Mur ligases lie within the central ATP-binding domain, those in the N- and C-termini generally do not co-localise with the known substrate binding residues.

Amino acids of published importance for ATP binding (species abbreviation subscripted); the P-loop within TGTXGKT^Sa^, E220^Mt^, D356^Sa^, N347^Mt^, R377^Mt^ and R392^Mt^ are conserved in the plant enzymes *M. endlicherianum* MurE and PpMurE, as well as a lysine, K219^Sa^, carbamylated in MurD for positioning the MgATP complex for the generation of a transient UDP-Mur*N*Ac-phosphodi-peptide intermediate (Dementin et al., 2001). K360^Sa^ and Y343^Mt^ have undergone conservative changes. Similarly, residues that bind UDP-Mur*N*Ac, S28^Ec^, HQA45^Ec^, NTT158^Ec^, E198^Mt^, S184^Ec^, QXR192^Ec^ and H248^Mt^ are no less conserved in plants than they are between bacteria.

Although most of the UDP-Mur*N*Ac-tripeptide interactions are within the MurE central domain, those made in relation to the appended amino acid, D,L-DAP or L-Lys, are within the C-terminal domain. All of the identified bacterial MurE residues that interact with D,L-DAP are highly conserved in plant MurE proteins. More specifically, with reference to *E. coli* MurE and *M. tuberculosis* MurE, it is possible to distinguish those that interact with either the D- or L-stereocentre carboxylates of D,L-DAP : G464^Ec^, E468^Ec^, D413^Ec^ and N414^Ec^, which bond to the D-stereocentre, R389^Ec^, which bonds with the L-stereocentre, and especially R416^Ec^, which interacts with both the L- and D-centre carboxylates. Of these R389^Ec^, N414^Ec^, R416^Ec^, G464^Ec^ and E468^Ec^ are less consistently present in MurE ligases from Gram positive bacteria that incorporate L-Lys, a decarboxylated derivative of D,L-DAP, which has only been reported to interact with the R383^Sa^, D406^Sa^ and E460^Sa^ residues (Ruane et al., 2013). Similarly, the pattern of charged residues in the C-terminal domain of the basal streptophyte MurE (those highlighted red or purple in Supplemental Figure S9) would indicate a binding cleft for the amino acid substrate that is more basic and resembles that of the Gram negative MurE ligases. Together these data are in complete accord with our kinetic findings that D,L-DAP is the preferred substrate in plants and AnMurE, rather than L-Lys. As would be anticipated from the phylogeny, the more closely related *G. bacterium* MurE aligns strongly with the Gram negative DL-DAP incorporating enzymes, and includes the DNPR motif, which confers specificity for the D-stereocentre carboxyl and amino groups of D,L-DAP, indicating that this phylum is most likely to incorporate DL-DAP.

## Discussion

### *P. patens* peptidoglycan is synthesized from a UDP-Mur*N*Ac-D,L-DAP-pentapeptide

Growth of *P. patens* on the antibiotics phosphomycin, D-cycloserine and ampicillin facilitated the detection, by mass spectrophotometric analysis of the TCA-extracted metabolome, of peptidoglycan intermediates up to UDP-Mur*N*Ac-Ala-Glu-DAP-Ala-Ala in the moss. These data enable us to conclude that the identical basic building blocks for the Gram negative bacterial cell wall are found in basal embryophytes. With evidence for knock-out phenotypes for *P.patens* homologs of bacterial *MraY*, *MurJ* and *PBP1A* and the presence of mRNA for *MurG* (Machida et al., 2006; Homi et al., 2009; Utsunomiya et al., 2020) it would be expected that the D,L-DAP-containing pentapeptide within the stroma is lipid-linked then flipped across the chloroplast inner envelope membrane and polymerised into peptidoglycan to form a ‘sacculus’ bounding the organelle, as indicated from fluorescent-labelling using a D-Ala-D-Ala analogue (Hirano et al., 2016). By analogy with bacteria and from the predicted transit peptides of the peptidoglycan-maturing proteins it is anticipated that the peptidoglycan will lie between the inner and outer membranes of the chloroplast, although this has yet to be determined (Figure 1).

### PpMurE appends D,L-DAP to UDPMurNAc-Ala-Glu

From our data, it is evident that the moss MurE ligase, with the transit peptide omitted, PpMurE_L63, can efficiently append D,L-DAP to UDP-Mur*N*Ac-L-Ala-D-Glu *in vitro*, as can the cyanobacterial enzyme from *Anabaena* sp. strain PCC 7120, AnMurE. This is in accordance with the D,L-DAP content of peptidoglycan in the cyanobacteria *Synechococcus* sp. and *Synechocystis* sp. (Jurgens et al., 1983; Woitzik, 1988) and is inconsistent with the observation that *Anabaena cylindrica* may incorporate L-Lys (Hoiczyk and Hansel, 2000). Our *in vitro* MurE enzymological data also complement the mass spectrometric analysis of the antibiotic-grown *P. patens* which identified UDP-Mur*N*Ac-D,L-DAP intermediates as being present *in vivo* in the TCA-extracted metabolome.

That UDP-Mur*N*Ac-L-Ala-D-Glu is an efficient substrate for PpMurE_L63 is significant in that there is no obvious homolog in most green plants for glutamate racemase (MurI), exceptions include the glaucophyte alga *Cyanophora paradoxa* (Contig25539), the charophyte alga *K. nitens* (GAQ85716.1) but not *M. endlicherianum*, a zygenematophycean alga proposed to be closest to the embryophyte branch point. Here, this function may be replaced by a D-alanine amino transferase (DAAA), of which there are two genes having weak homology to *Bacillus subtilis* DAAA in both *P. patens* and *M. endlicherianum* (Phytozome v.13 *P. patens*: Pp3c6_5420 (15.7%), Pp3c16_17790 (14.7%) and OneKP *M. endlicherianum*: WDCW scaffolds 2009723 (17.6%) and 2007189 (16.5%)). Alternatively *P. patens* diaminopimelate epimerase (DapF), like Chlamydial DapF, may possess the dual specificity required to racemase L-Glu to D-Glu in addition to its epimerization of L,L-DAP to D,L-DAP (De Benedetti et al., 2014).

### Substrate Preference of AnMurE and PpMurE

The high degree of specificity of both AnMurE and PpMurE_L63 for D,L-DAP, over the alternatives L,L-DAP, D,D-DAP and L-Lys, is consistent with other D,L-DAP-incorporating enzymes assayed *in vitro*, including *E. coli* MurE, *M. tuberculosis* MurE and *Chlamydia trachomatis* MurE, for which L-Lys is either a very poor substrate or is not accepted at all (Supplemental Table S7). Similarly, the L-Lys-incorporating *S. aureus* MurE does not incorporate D,L-DAP *in vitro*. Not all MurE ligases are as selective, *Thermotoga maritima* MurE incorporates L-Lys and D-Lys in almost equal amounts *in vivo* (Huber, 1986) and can efficiently incorporate D,L-DAP *in vitro* (Boniface et al., 2006). In this regard it is notable that *T. maritima* MurE possesses a DDP**R** motif, which includes the arginine residue of the consensus DNPR of D,L-DAP-incorporating enzymes which hydrogen bonds to and stabilises D,L-DAP, consequently the almost complete absence of D,L-DAP in *T. maritima* peptidoglycan has been attributed to its low intracellular concentration. This almost absolute specificity of most MurE ligases is indicative of a requirement that the stem peptide be composed of the correct amino acids to facilitate optimal transpeptidation (Vollmer et al., 2008).

### PpMurE is a slow but efficient MurE ligase

Kinetic analyses of PpMurE_L63 demonstrated an enzymatic efficiency similar to bacterial MurE homologs, as estimated by comparison of K_cat_^App^/K_M_^App^ (Supplemental Table S7). Further comparisons with other D,L-DAP-incorporating enzymes, and in particular those of the obligate intracellular pathogens *C. trachomatis* and *M. tuberculosus*, revealed the plant MurE to have a similarly low K_M_ for the amino acid substrate relative to the L-Lys-incorporating enzymes. This may reflect either (or both) a lower abundance of D,L-DAP or the potential toxicity of the D,L-diamino acid, particularly in a eucaryotic cell (Kolukisaoglu and Suarez, 2017). A higher K_M_ for L-Lys-incorporating MurE ligases has been attributed to the much greater abundance of this amino acid in bacteria (Mengin-Lecreulx et al., 1982; Ruane et al., 2013). The availability of the D,L-DAP substrate in plants, as in cyanobacteria and Chlamydiae, is not in question as the biosynthesis of L-Lys is catalysed by DAP decarboxylase (LysA) from D,L-DAP which is ultimately derived from aspartate (Hudson et al., 2006).

Comparison of the PpMurE_L63 K_cat_^App^ with the bacterial enzymes reveals the rate of turnover to be quite low, possibly reflecting the apparent low density of peptidoglycan surrounding the chloroplast and a concomitant slower rate of synthesis compared to rapidly dividing, free-living bacteria. Moreover, the plant enzyme has UDP-Mur*N*Ac-Ala-Glu kinetics best fitted to a substrate inhibition model, possibly to ensure that peptidoglycan synthesis proceeds at a rate insufficient to consume the majority of available prenyl phosphates that are otherwise required for other pathways.

It is important to mention that the *P. patens* genome encodes two MurE homologs (PpMurE1: Pp3c23_15810, studied in this paper, and PpMurE2: Pp3c24_18820) which have 72.2% amino acid identity to each other over the conventional bacterial MurE ligase domains and 48.4% identity overall. PpMurE2 primarily differs from PpMurE1 in encoding a long, relatively unstructured extension at the amino terminus and a short carboxy terminal extension (expanded description in Supplemental Figure S10). Although the DNPR motif and other amino acids associated with D,L-DAP binding are retained in PpMurE2, knock out mutations of PpMurE1 alone results in a comprehensive macrochloroplast phenotype (Machida et al., 2006; Garcia et al., 2008), consistent with the hypothesis that this protein is sufficient for peptidoglycan synthesis in the moss. Moreover, preliminary *in vitro* experiments indicate that intact PpMurE2 does not function as a MurE ligase (data not shown) and we would suggest that both the amino and carboxy terminal extensions have been acquired during streptophyte evolution to participate in novel interactions thereby facilitating an alternative function for MurE within the chloroplast transcription and translation apparatus.

In contrast to *P. patens* (and the Polypodiidae ferns, Supplemental Figure S10), many in the same and closely related phylla encode a single *MurE* homolog with both the amino and carboxy terminal extensions and aligning more closely to PpMurE2, yet these proteins would be anticipated to function as MurE ligases. We propose the shorter MurE in *P. patens* and the Polypodiidae ferns represents a de-evolution of streptophyte MurE to more closely resemble its bacterial counterpart. It has yet to be determined at what point in streptophyte evolution the function of MurE changed and whether in any plants it remains a bifunctional protein capable of both MurE ligase activity and interaction with chloroplast RNA polymerase in chloroplast transcription.

That basal embryophyte MurE has evolved a new role essential to plastid photomorphogenesis in seed plants indicates an exaptation from its original function in peptidoglycan biosynthesis and plastid division (Williams-Carrier et al., 2014). This raises the intriguing question why important residues of the D,L-DAP-binding motif are retained, in similar proximity to the ATPase domain, in these proteins. We would speculate that the novel function of the MurE-like proteins in seed plants could have evolved consequent on the two whole gene duplication events which occurred in an ancestral moss, as opposed to in the liverworts or hornworts (Lang et al., 2018).

### Predicted streptophyte peptidoglycan structure from peptidoglycan gene homologies

The moss ‘sacculus’, like that of Chlamydiae, has been recalcitrant not only to visualisation by electron microscopy but also to common extraction protocols, making analysis of the mature polymer a future goal. The moss chloroplast envelope membranes were found to be closely appended with little dense intervening material (Takano and Takechi, 2010; Matsumoto et al., 2012; Sato et al., 2017), likewise in Chlamydiae the apparent deficit of a bounding sacculus lead to the term the ‘chlamydial anomaly’ (Packiam et al., 2015). This is in marked contrast to most cyanobacteria where the cell wall is highly cross-linked and forms a broad, electron dense layer (Hoiczyk and Hansel, 2000). Intermediate between these extremes is the earliest side branch in plant evolution, the glaucophyte algae, where the cyanelles comprise a peptidoglycan layer that has been more tractable to visualisation and analysis (Pfanzagl et al., 1996; Higuchi et al., 2016).

It would appear that progressive transition of a bacterium from free-living to endosymbiont or pathogen and thence to an integrated organelle is associated with a reduction in substance of the sacculus. Presumably there are not the same osmotic constraints and risks of dehydration within the host cell and the vestigial peptidoglycan may function primarily or exclusively for the purpose of assembly of the division apparatus. Additionally, it may be that for cyanobacterial evolution into a cyanelle and subsequently a plastid that a finer, net-like cell wall would be a prerequisite if extensive exchange of larger molecules, including lipids and proteins, were to occur. Supportive of this suggestion is the fact that most of the bacterial PBPs which cross-link the lipid-linked Glc*N*Ac-Mur*N*Ac-pentapeptide precursor, have been identified as having no predicted product from RNA-seq data (data not shown). Currently the only reported exception is a PBP1A homolog, the transpeptidase and transglycosylase functions of which have an almost complete knock out phenotype (Machida et al., 2006; Takahashi et al., 2016).

We also propose that streptophyte peptidoglycan must differ in its mature form by being uniquely modified to distinguish it from the peptidoglycan of potential plant pathogens. The *P. patens* genome encodes a battery of proteins that include peptidoglycan-binding and LysM domains and which frequently but not invariably include cell export signals (data not shown). Many of these proteins will be part of the defences of the plant cell which are activated on detection of fungal and bacterial cell wall material. To evade the host cell defences it is anticipated that an endosymbiont, obligate pathogen or evolving organelle must protect its peptidoglycan from the host defences, conceivably by modification of the peptide stem (Wolfert et al., 2007) or the Glc*N*Ac-Mur*N*Ac backbone (Davis and Weiser, 2011). Predictions as to what those modifications might be in streptophytes are hampered by the fact that the ancestry of the modifying enzymes is not necessarily cyanobacterial. We have reported here the closer homology of PpMurE to MurE in the Gemmatimonadetes phylum and we can further include *P. patens* PBP1A, MurF, MurD, MurG and Ddl as most closely related to homologs within the same Fibrobacteres-Chlorobi-Bacteroidetes group of Gram negative bacteria (data not shown). The diverse origins of several peptidoglycan biosynthesis-related proteins have previously been reported (Sato and Takano, 2017). Therefore, it appears highly probable that a horizontal gene transfer event of a distinct Gram negative peptidoglycan-related gene cluster must have occurred early in the plant lineage. Hence we conjecture a simultaneous transfer of peptidoglycan-modifying genes could have occurred that would introduce novel modifications to the mature polymer, distinct from any in cyanobacteria. This is not without precedent, as the divergent glaucophyte algae were found to append *N*-acetyl-putrescine to the second residue in the stem peptide (Pfanzagl et al., 1996).

Here we have determined that chloroplast peptidoglycan in the streptophyte, *P. patens*, is constructed from typical Gram negative UDP-MurNAc-D,L-DAP-pentapeptide peptidoglycan precursor. However, we propose that the final polymerised structure derived from this building block differs from its cyanobacterial progenitor by being both less highly polymerised and, to distinguish it from plant pathogens and thereby evade the plant immune response, significantly modified.

## Supplemental Data

Supplemental Text S1. Effects of antibiotics on *P. patens*

Supplemental Figure S2. Negative ion nanospray TOF mass spectra of TCA-extracted peptidoglycan intermediates

Supplemental Figure S3. PAGE gel of AnMurE and PpMurE_L63 after gel filtration

Supplemental Figure S4. D,L-DAP substrate curves for AnMurE and PpMurE_L63

Supplemental Figure S5. Assay data demonstrating PaMurF utilises the product of AnMurE and PpMurE_L63

Supplemental Text S6. Activities of AnMurE and PpMurE_L63 with pH and buffer

Supplemental Table S7. Comparison of AnMurE and PpMurE_L63 kinetics with published data for other MurE ligases

Supplemental Figure S8. Neighbour joining phylogram of MurE of Gram-negative bacteri and early plant species

Supplemental Figure S9. Clustal Omega multiple sequence alignment of MurE homologs

Supplemental Figure S10. Phylogram of evolutionary relationship of both PpMurE proteins to selected MurE homologs

## Acknowledgements

The authors thank Professor Hiroyoshi Takano (Kumamoto University, Japan) for kindly providing the pTFH22.4 vectors with *Anabaena* (PCC7120 Q8YWF0|MURE_NOSS1) and *P.patens* (Pp3c23_15810V3.2) *MurE* cDNA and Dr Sven Gould (Heinrich Heine University, Düsseldorf, Germany) for helpful discussion on chloroplast evolution. We are also grateful to Julie Tod and Anita Catherwood (University of Warwick, UK) for synthesis of UPD-Mur*N*Ac-dipeptide, - tripeptide and -pentapeptide and providing *Streptococcus pneumoniae* MurE and *Pseudomonas aeruginosa* MurF and Ian Hands-Portman for access to and training in the School of Life Sciences Imaging Suite, University of Warwick, UK). We also gratefully acknowledge Prof. Rebecca Goss (St Andrews, UK) for provision of pacidamycin.

---

**Supplemental Text S1** Effects of antibiotics on *P. patens*

Antibiotics which only rarely effected macrochloroplast development included vancomycin (1.0, 5.0 and 25 µg.ml-1), a 1.45 kDa glycopeptide which binds D-Ala-D-Ala, thereby inhibiting PBP transpeptidation, and the 1.42 kDa bacitracin (20 and 100 µg.ml-1), which complexes with C55-isoprenyl pyrophosphate, inhibiting recycling (Figure 1^8,6^ and Figure 2, C and E). Neither antibiotic typically traverse cytoplasmic membranes, and therefore a strong phenotype was not expected as it is anticipated they would have to penetrate not only the cytoplasmic membrane but also, potentially, the outer chloroplast membrane. It may be that any observed effect of these antibiotics was restricted to damaged or senescing cells. At high concentrations (500 µg.ml-1) bacitracin did cause premature senescence.

A22 hydrochloride, a smaller molecule at 271.6 kDa, was tested at 2.5 and 10 µg.ml-1 and was likewise found to result in macrochloroplast formation in some but not most cells, although at higher concentrations its impact was more pleiotropic and chloroplasts were considerably bleached (Figure 2, H and I). A22 inhibits MreB, an actin homolog and cytoskeletal protein that controls bacterial width in rod-shaped bacteria by spatiotemporal regulation of peptidoglycan synthesis. Since there is not an evident MreB homolog in the moss (Ozdemir et al., 2018), any effect of A22 may be consequent on a less specific effect on chloroplast heat shock proteins having homology to MreB, especially HSP70 (Gao and Gao, 2011).

Another antibiotic clearly pleiotropic in its effect was tunicamycin (0.2, 1.0 and 5.0 µg.ml-1), a glycoprotein that inhibits the transfer of phospho-MurNAc-pentapeptide to the lipid carrier undecaprenyl pyrophosphate by MraY (Figure 1^4^). At concentrations equal to or above 1 µg.ml-1 it caused chloroplast malformation, slow growth and apoptosis (data not shown). This could be attributed to its effect on the maturation of glycoproteins in the endoplasmic reticulum since, in eucaryotes, tunicamycin also blocks the transfer of UDP-GlcNAc to dolichol phosphate.

Pacidamycins 1 and 5, cationic peptides with homology to the bacteriophage øX174 lysis protein Arg-Trp-x-x-Trp motif, believed to bind the cytoplasmic surface of MraY and thereby inhibiting it (Figure 1^3^) (Rodolis et al., 2014; Bugg and Kerr, 2019), had little effect on either growth rate or chloroplast division (data not shown). Likewise, Murgocil, a 448Da steroid-like molecule, which inhibits peptidoglycan synthesis in *Staphylococcus aureus* and is predicted to bind in the MurG active site blocking UDP-GlcNAc access (Figure 1^5^), when tested at 1, 5 and 25 µg.ml-1 was found to have little effect on protonemata phenotype (Figure 2, F).

The effect of the three antibiotics, phosphomycin, D-cycloserine and ampicillin (Figure 3 B,D and G), subsequently selected for investigating the accumulation of peptidoglycan intermediates is detailed in the text of the paper.

**Supplemental Figure S2.**
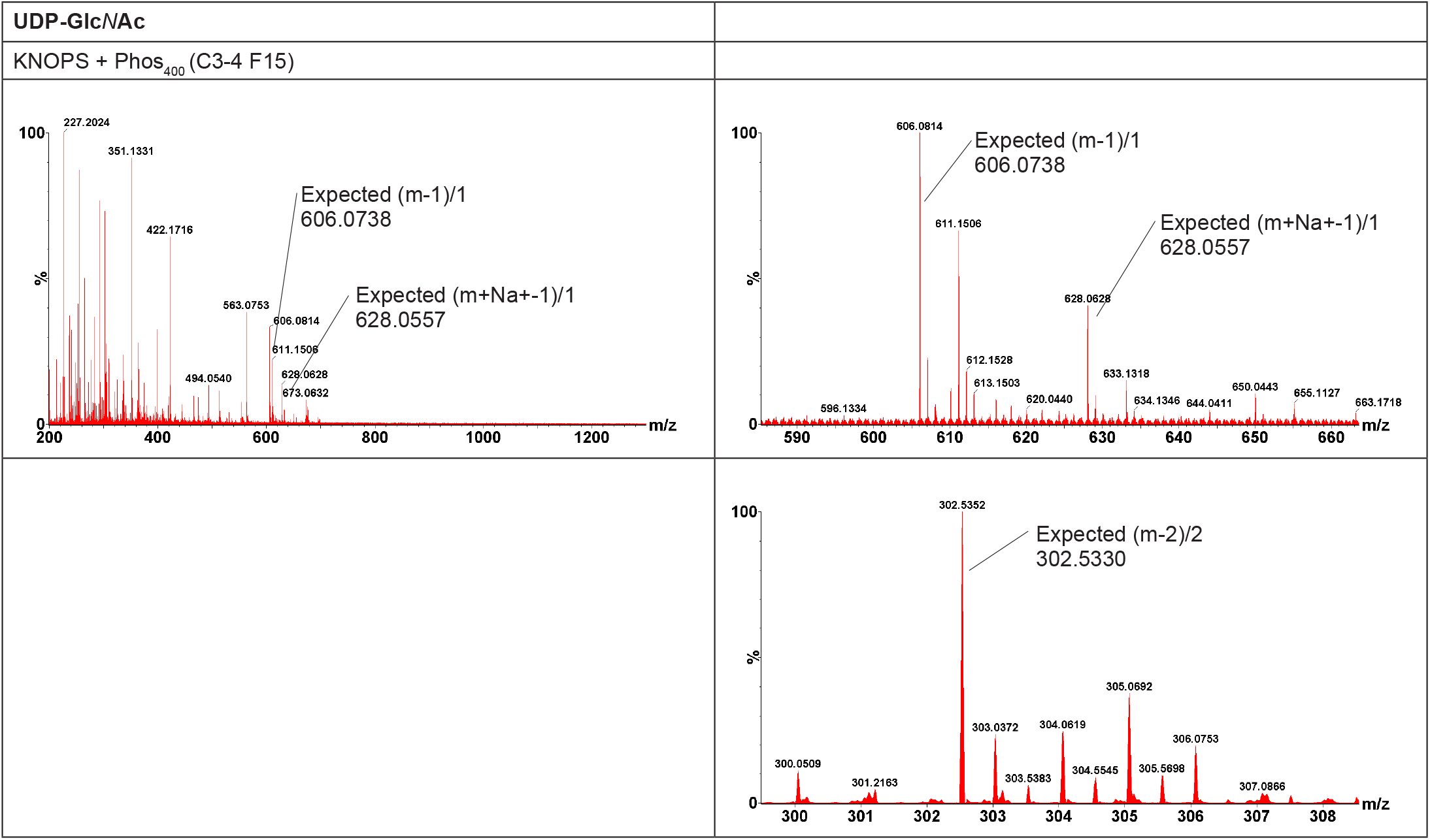
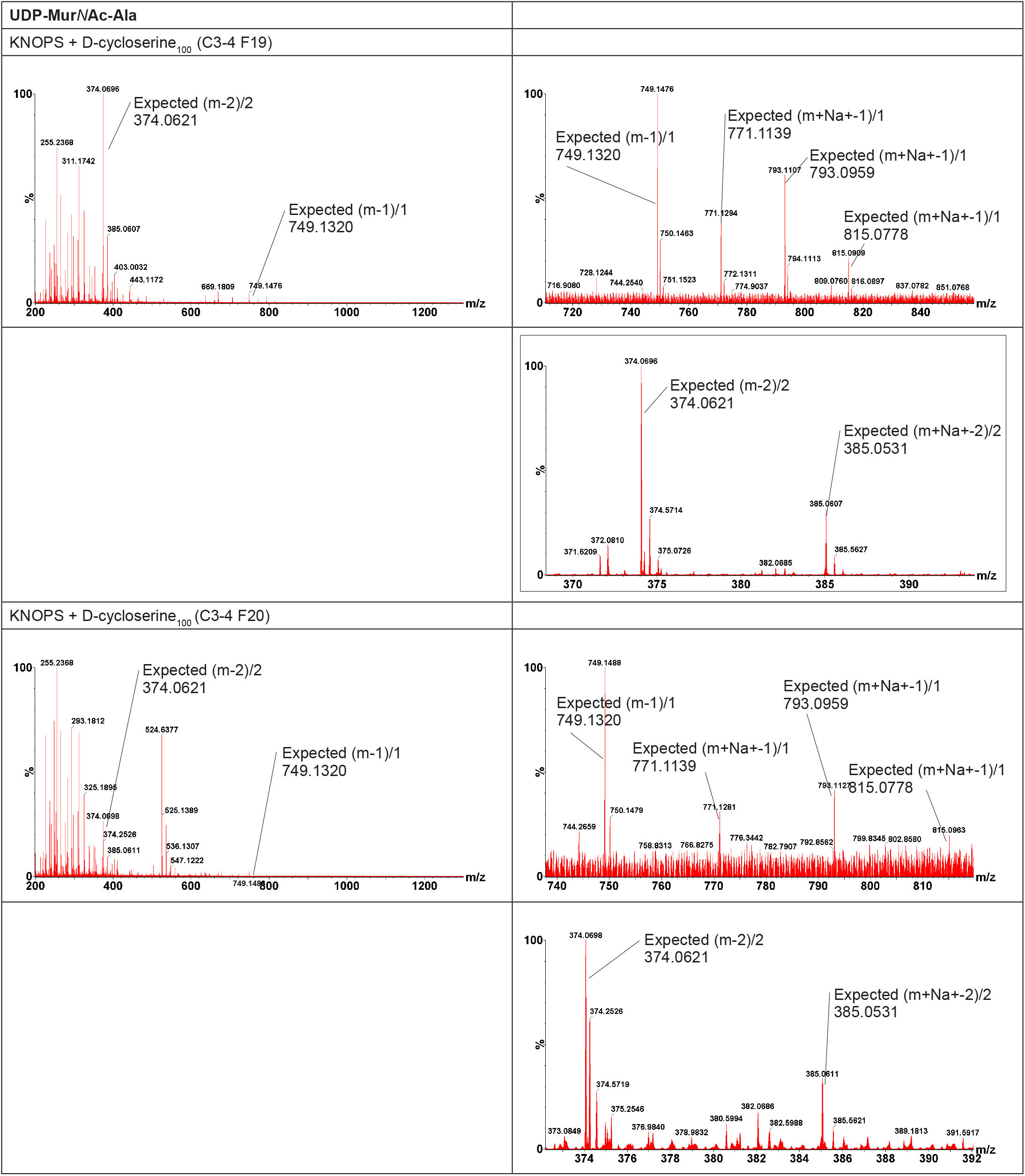
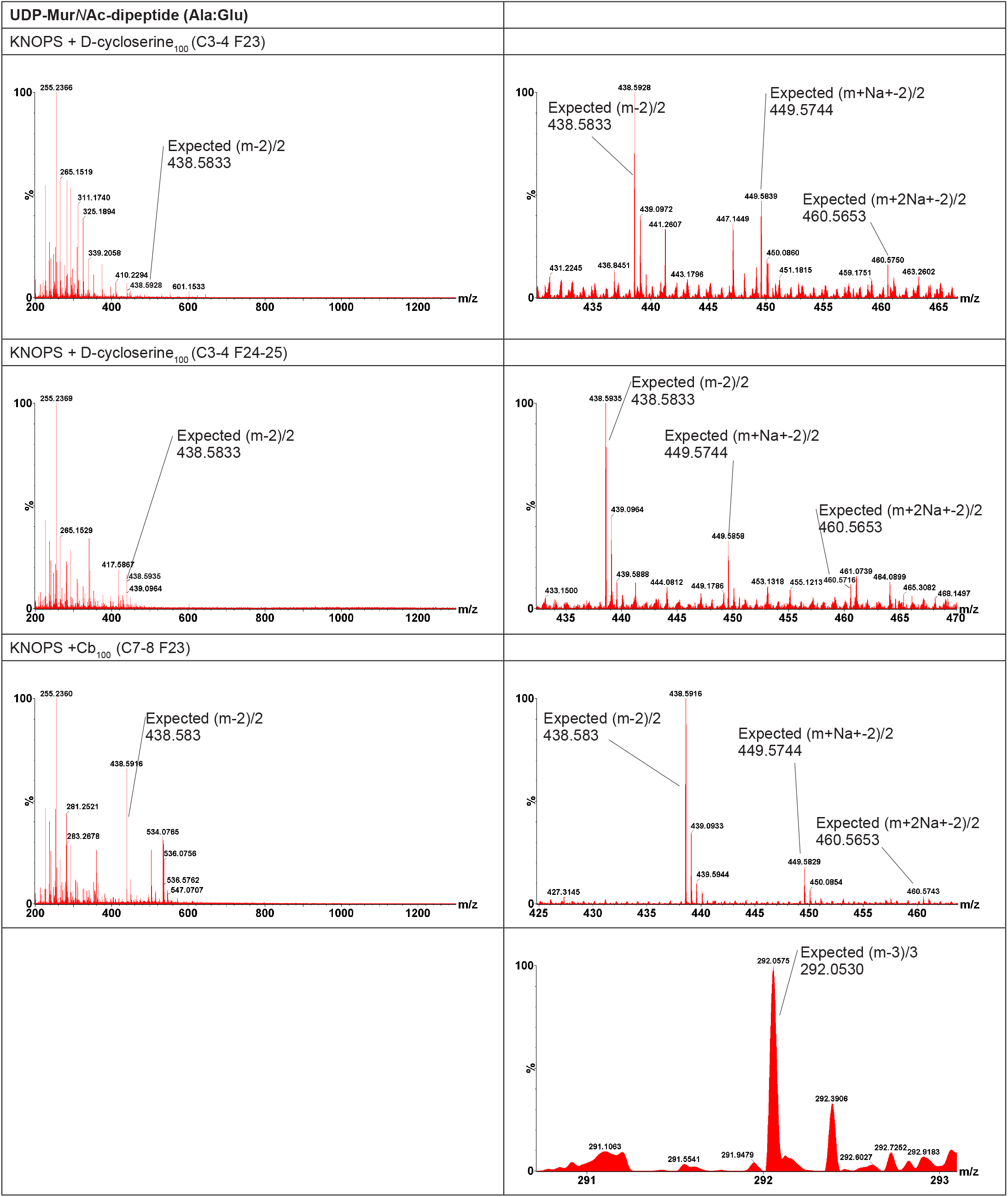
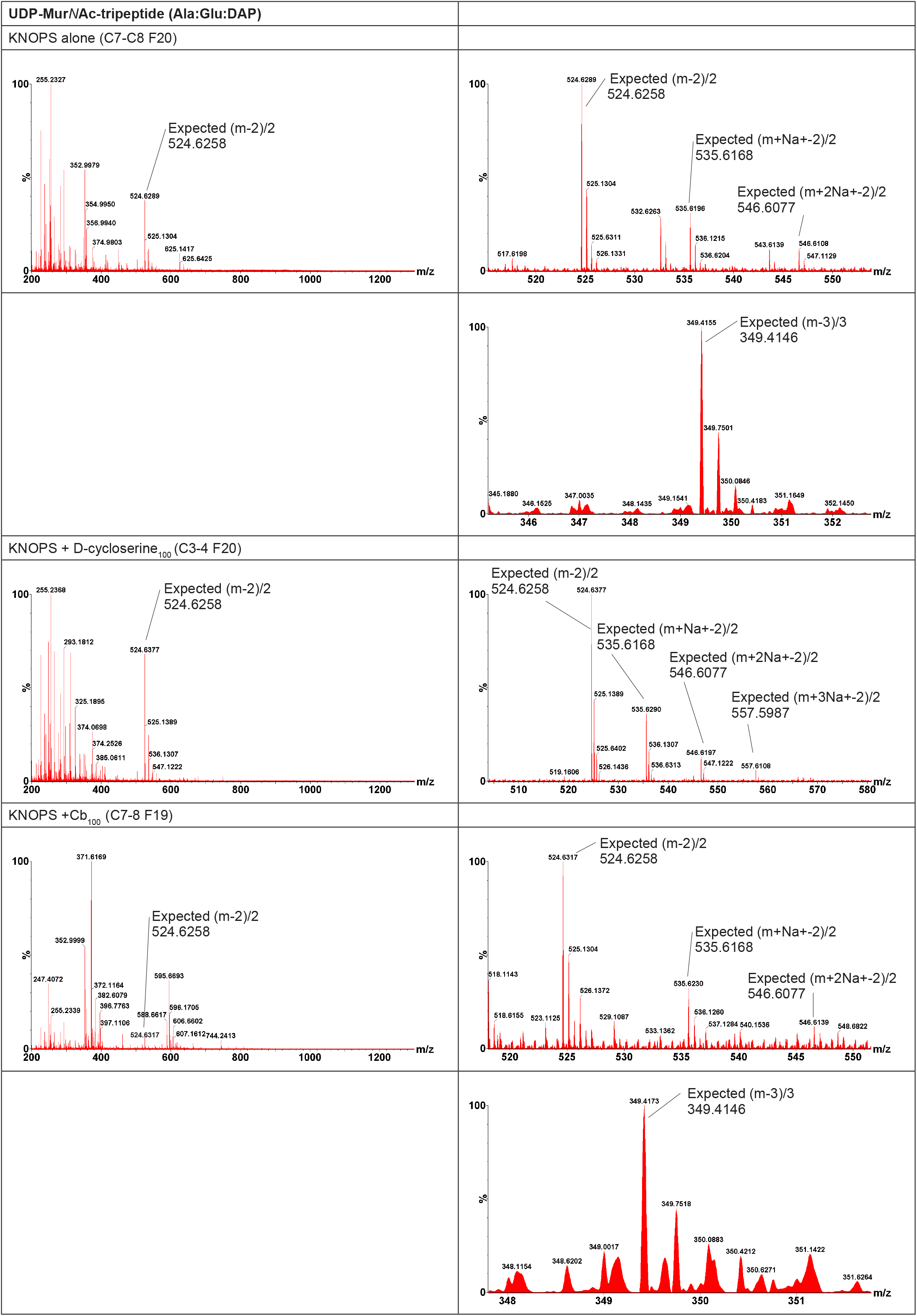
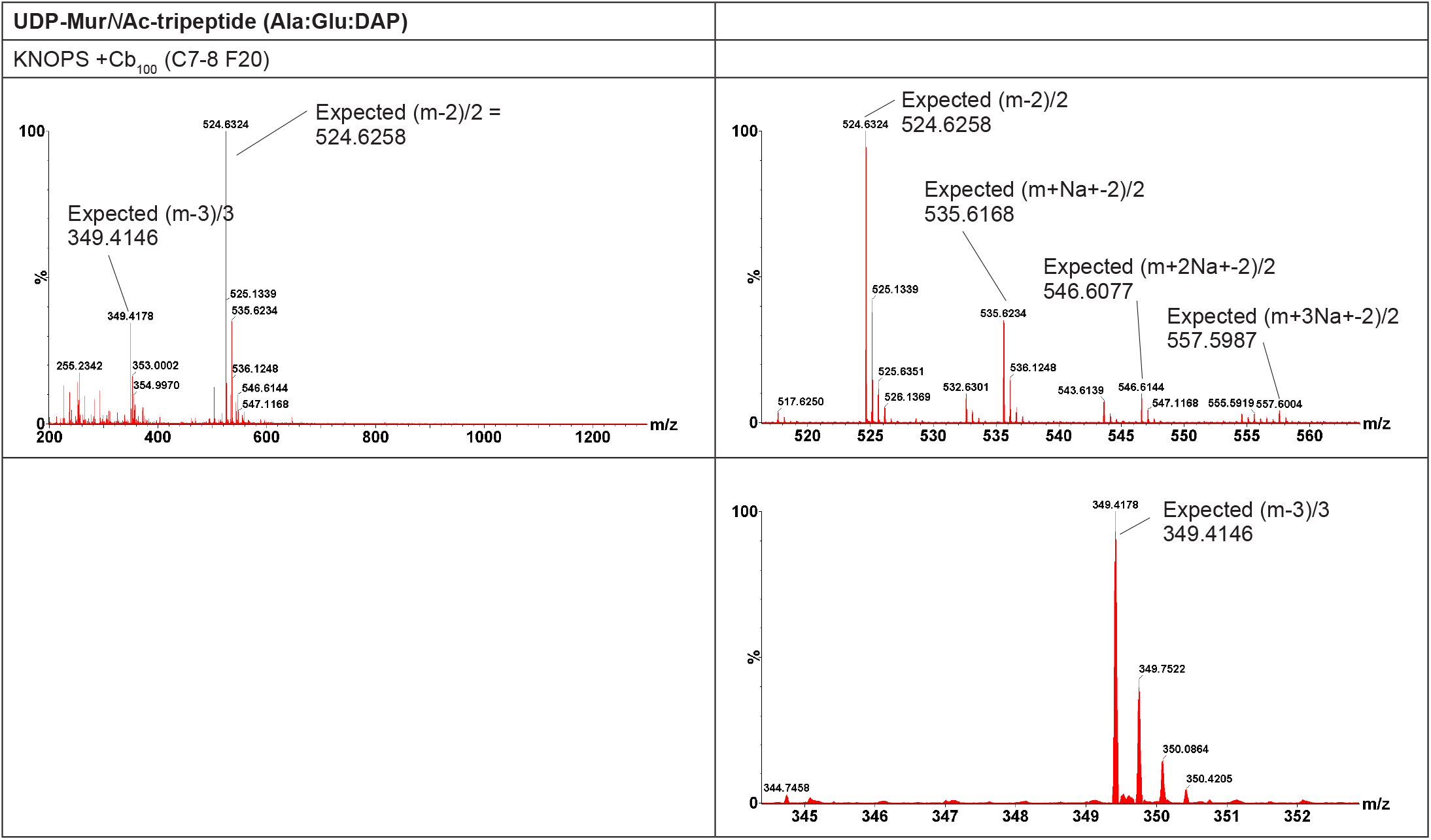
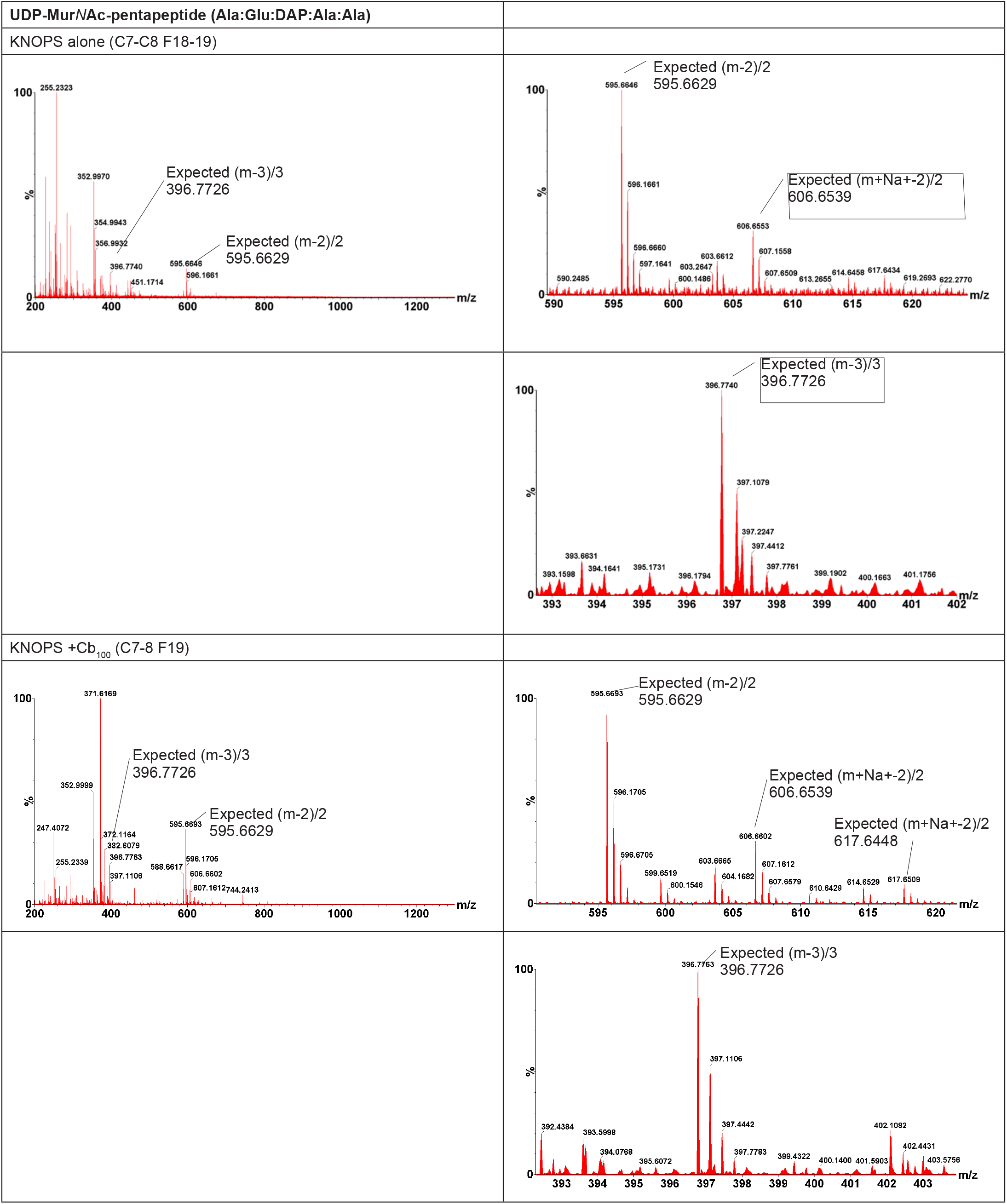
Negative ion nanospray TOF mass spectra of TCA-extracted peptidoglycan intermediates, with the expected mass:charge (mz) values for the different species in red boxes. *P. patens* was grown on KNOPS medium with and without antibiotics, including Phos_400_ (phosphomycin 400 μg.ml^-1^), D-cycloserine_100_ (D-cycloserine 100 μg.ml-1) and Cb_100_ (carbenicillin 100 μg.ml-1). UDP-linked intermediates were purified by chromatography on Superdex Peptide and then MonoQ columns and their respective fractions (C and F) are indicated in brackets in the headers.

**Supplemental Figure S3.**
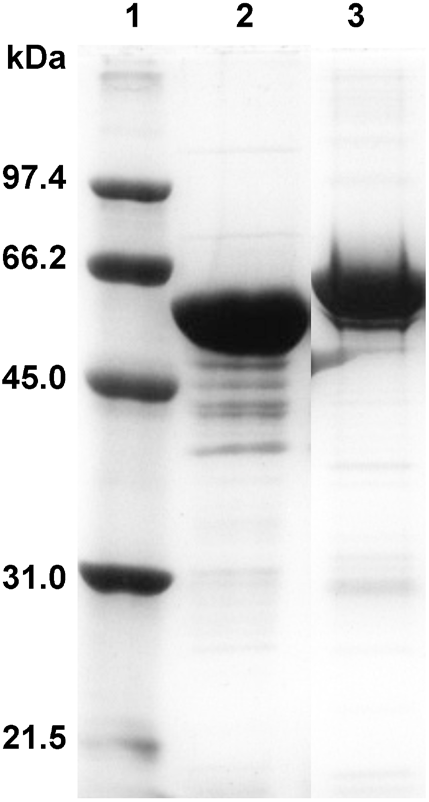
PAGE gel of AnMurE and PpMurE_L63 after gel filtration. Lane 1, protein size marker, 2, AnMurE (predicted mass 56.57 kDa) and 3, PpMurE_L63 (predicted mass 62.59 kDa).

**Supplemental Figure 4.**
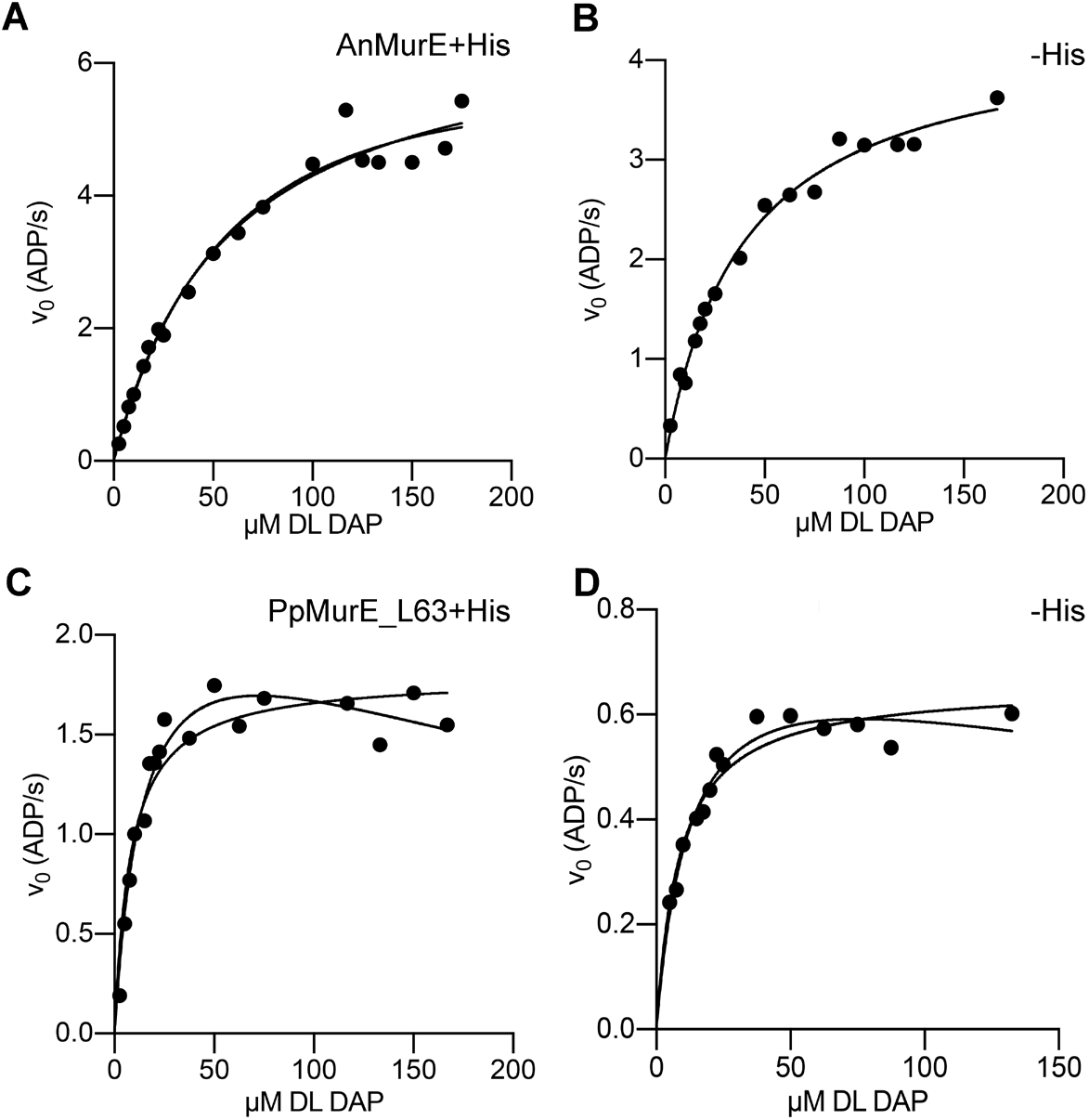
D,L-DAP substrate curves for AnMurE (A,B) and PpMurE_L63 (C,D). A and C, substrate curves for AnMurE and PpMurE with His tags, at 50 nM and 146 nM, respectively. Band D, substrate curves for AnMurE and PpMurE_L63 after His tags have been cleaved by TEV protease, at 76 nM and 328 nM respectively. Assays were with 1 mM UDP-MurNAc-dipeptide in 50 mM PIPES pH 6.7 (AnMurE) or 50 mM Tricine pH 8.7 (PpMurE_L63). Rates (v_0_) in ADP.s·^1^ are mols ADP.mol Mur ligase-^1^.s-^1^. Data show Michaelis Menten curves superimposed on those for substrate inhibition and indicate best fit to Michaelis Menten kinetics for AnMurE and to substrate inhibition for PpMurE_L63 (R^2^ values for Michaelis Menten and substrate inhibition are in Figure 4, B).

**Supplemental Figure S5.**
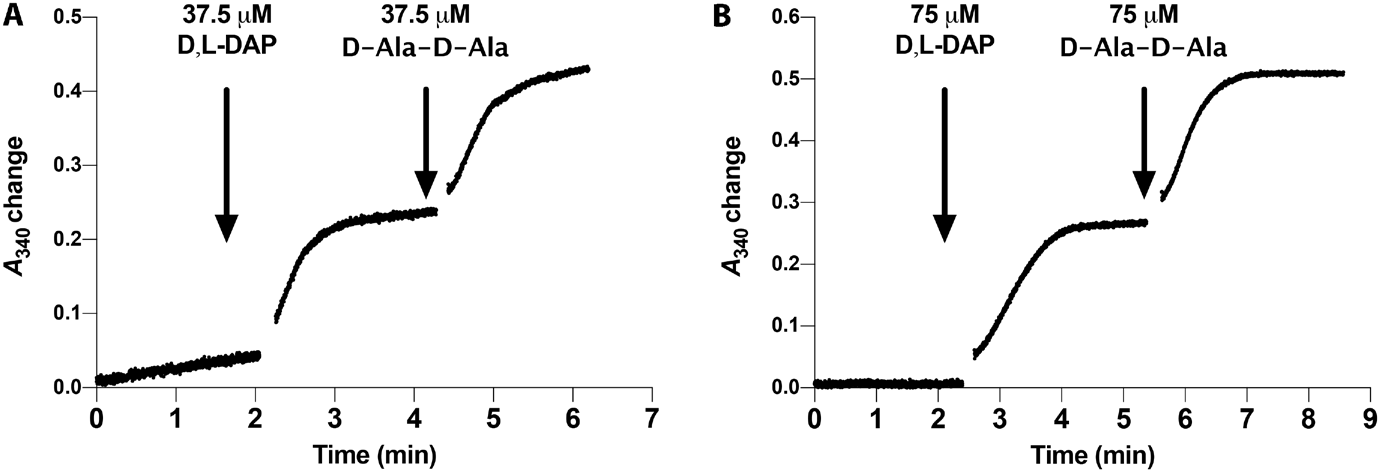
Assay data demonstrating PaMurF utilises the product of AnMurE and PpMurE_L63. A, AnMurE and B, PpMurE_L63. Change in NADH absorbance at A 340 is coupled to the release of ADP by the MurE and MurF ligases on addition of their substrates D,L-DAP and D-Ala--D-Ala, respectively. Assays included 492 nM PaMurF in 50 mM Hepes, pH 7.6, 375 µM UDP-MurNAc-dipeptide and A, 100 AnMurE+His or B, 300 nM PpMurE_L63+His.

**Supplemental Figure S6.**
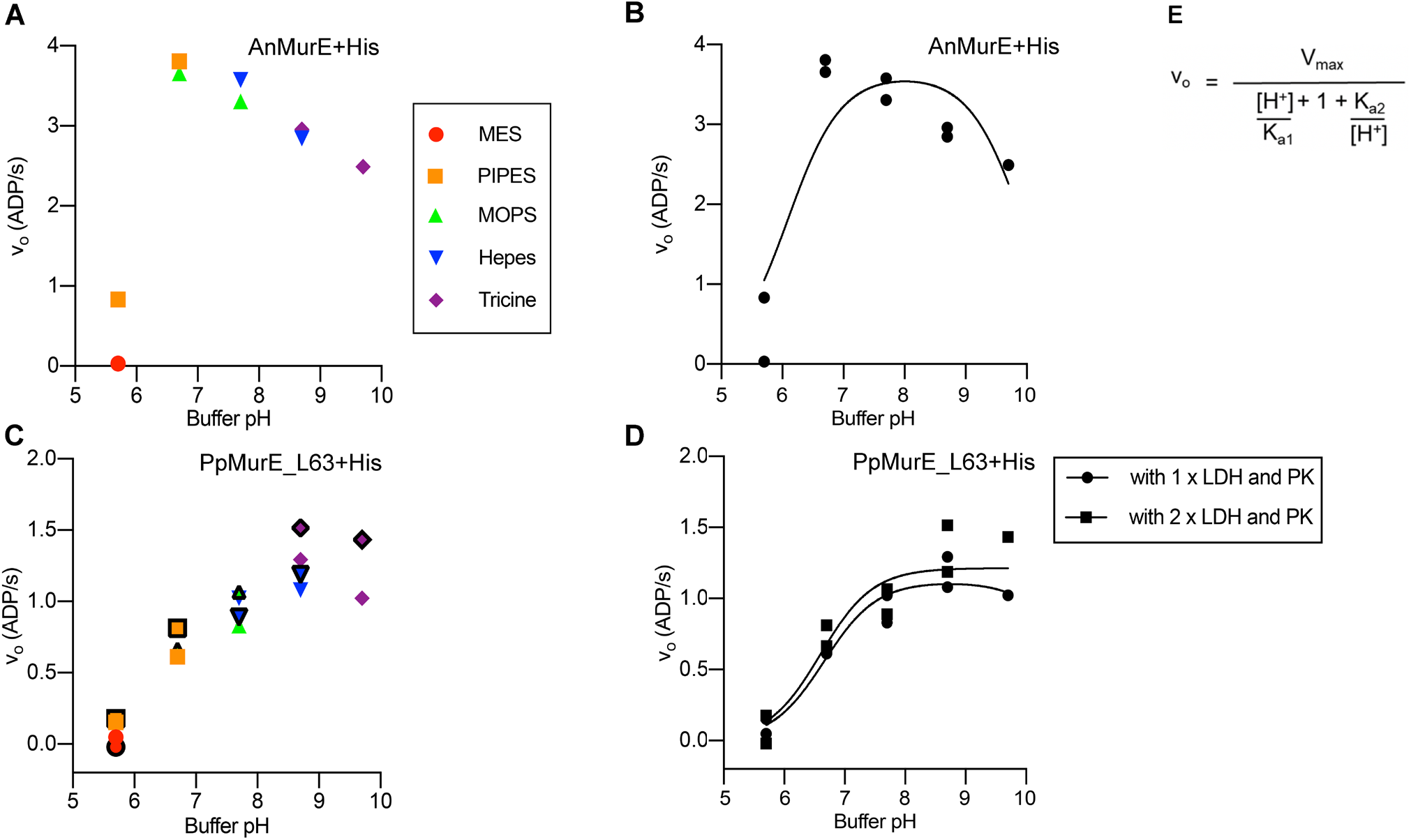
Activities of AnMurE and PpMurE_L63 with pH and buffer. Data are presented in two ways: A and C, with coloured symbols to indicate buffers (see legend) and B and D, with non-linear fit to estimate pH optima.. Assays for A and B, AnMurE+His and C and D, PpMurE_L63+His, respectively, were in 50 mM buffers in the pH range 5.7-9.7. For PpMurE_L63 C, symbols with black outlines and D, square symbols represent assays with the coupling enzymes lactate dehydrogenase (LOH) and pyruvate kinase (PK) at double the normal concentration (see materials and method), to confirm these were not limiting. Assays included 260 µM UDPMurNAc-dipeptide and 150 µM D,L-DAP. If we make the assumption that the only variable responsible for a change in enzyme activity over the pH range tested is the change in [H+J we can derive an equation that follows the relationship of activity *versus* pH (E) where Ka_1_ and Ka_2_ are dissociation constants of ionizable groups responsible for the ascending and descending limbs of the pH profile. Data indicate that the pH optima for AnMurE and PpMurE_L63 are 7.5 and 7.5-8.5 respectively.

**Supplemental Table S7.**
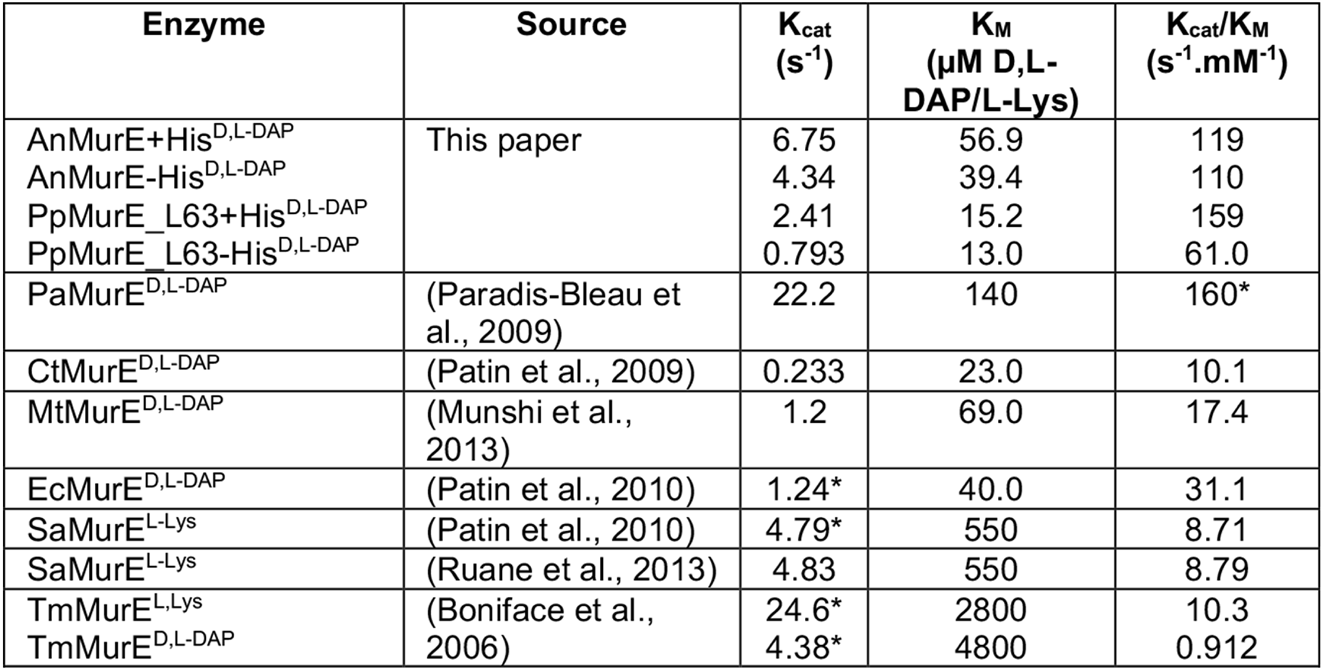
Comparison of AnMurE and PpMurE_L63 kinetics with published data for other MurE ligases. PaMurE^D,L-DAP^ *Pseudomonas aeruginosa,* CtMurE^D,L-DAP^ *Chlamydia trachomatis,* MtMurE^D,L-DAP^ *Mycobacterium tuberculosis,* E cMurE^D,L-DAP^ *Escherichia coli,* SaMurE^L-Lys^, *Staphylococcus aureus* and TmMurE^L-Lys^, *Thermotoga maritima.* Asterisks indicate where data were extrapolated from the published values.

| Enzyme | Source | $K_{cat}$<br>(s<sup>-1</sup>) | $K_M$<br>(μM D,L-DAP/L-Lys) | $K_{cat}/K_M$<br>(s<sup>-1</sup>.mM<sup>-1</sup>) |
| --- | --- | --- | --- | --- |
| AnMurE+His<sup>D,L-DAP</sup> | This paper | 6.75 | 56.9 | 119 |
| AnMurE-His<sup>D,L-DAP</sup> |  | 4.34 | 39.4 | 110 |
| PpMurE_L63+His<sup>D,L-DAP</sup> |  | 2.41 | 15.2 | 159 |
| PpMurE_L63-His<sup>D,L-DAP</sup> |  | 0.793 | 13.0 | 61.0 |
| PaMurE<sup>D,L-DAP</sup> | (Paradis-Bleau et al., 2009) | 22.2 | 140 | 160* |
| CtMurE<sup>D,L-DAP</sup> | (Patin et al., 2009) | 0.233 | 23.0 | 10.1 |
| MtMurE<sup>D,L-DAP</sup> | (Munshi et al., 2013) | 1.2 | 69.0 | 17.4 |
| EcMurE<sup>D,L-DAP</sup> | (Patin et al., 2010) | 1.24* | 40.0 | 31.1 |
| SaMurE<sup>L-Lys</sup> | (Patin et al., 2010) | 4.79* | 550 | 8.71 |
| SaMurE<sup>L-Lys</sup> | (Ruane et al., 2013) | 4.83 | 550 | 8.79 |
| TmMurE<sup>L-Lys</sup> | (Boniface et al., 2006) | 24.6* | 2800 | 10.3 |
| TmMurE<sup>D,L-DAP</sup> |  | 4.38* | 4800 | 0.912 |

**Supplemental Figure S8.**
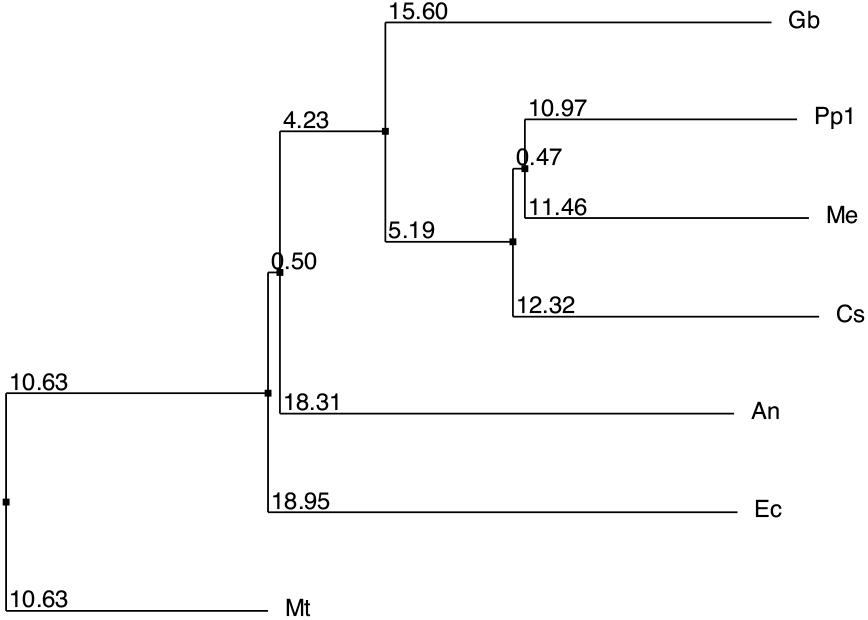
Neighbour joining phylogram of MurE of Gram negative bacteria and early plant species, computed using percentage identity in Jalview. Ec *E. coli* (strain K12), An *Anabaena nostoc* PCC7120, Mt *Mycobacterium tuberculosum*, Pp1 *P. patens* (Pp3c24_18820V3.2 v3.3 from Phytozome), Me *Mesotaenium endlicherianum* (WDCW from Onekp CNGBDB), Cs Coleochaete scutata (VQBJ from Onekp), Gb *Gemmatimonadetes bacterium*. All sequences are from the Uniprot or NCBI databases unless stated otherwise.

**Supplemental Figure S9.**
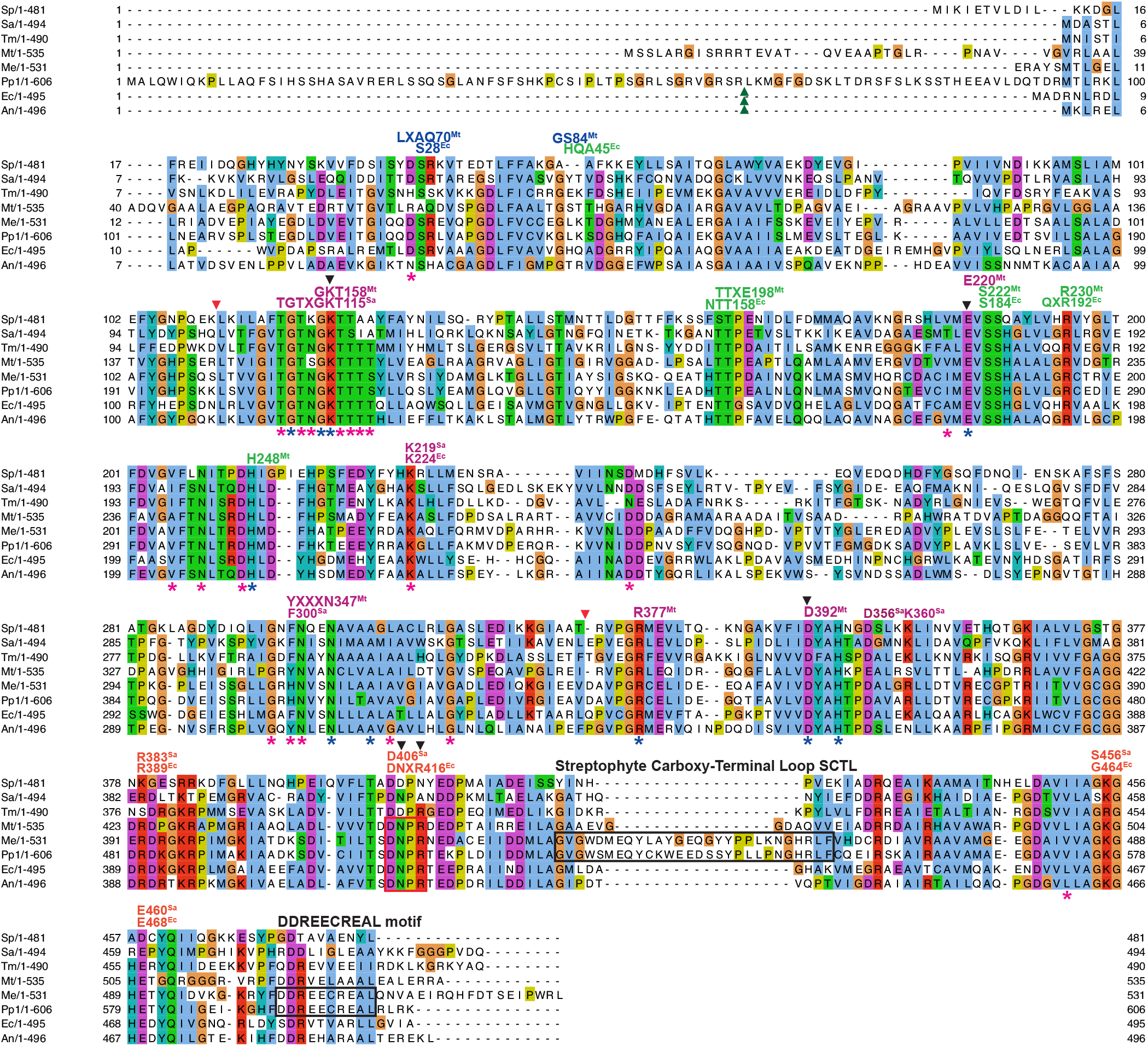
Clustal Omega (EMBL-EBI)(Madeira et al., 2019) multiple sequence alignment of MurE homologs displayed using Jalview (Waterhouse et al., 2009) with Clustalx designated colours: Sp *Streptococcus pneumoniae*, Sa *Staphylococcus aureus*, Mt *Mycobacterium tuberculosum*, Pp1 *P. patens* (Pp3c24_18820V3.2 v3.3 from Phytozome), Tm *Thermotoga maritima,* Me *Mesotaenium endlicherianum* (WDCW from Onekp CNGBDB), Ec *E. coli* (strain K12), An *Anabaena nostoc* PCC7120. All sequences are from the Uniprot or NCBI databases unless stated otherwise. Green arrows indicate ChloroP predicted cleavage site for PpMurE and red arrows the domain hinge points (Smith, 2010). Black arrows indicate residues with a reported role in MtMurE catalysis (Basavannacharya et al 2010). Letter labels indicate numbered residues with published ligand interractions: ^Ec^ for EcMurE (Gordon et al., 2001), ^Mt^ for MtMurE (Basavannacharya et al., 2010; Maitra et al., 2019) and ^Sa^ for SaMurE (Ruane et al., 2013) with colours indicating binding to UDP (blue), Mur*N*Ac sugar (green), ATP or ADP (mauve) and DL-DAP (orange) ligands. Blue asterisks indicate residues common to the Mur ligase family, which includes folylpolyglutamate synthetase, cyanophycin synthetase and the capB enzyme from Bacillales (Gordon et al., 2001; Smith, 2010) and pink asterisks indicate residues common to MurC, D, E and F ligases (Basavvanacharya et al., 2010). Two streptophyte-specific features are identified by black boxes and the DNPR consensus by a red box.

**Supplemental Figure S10.**
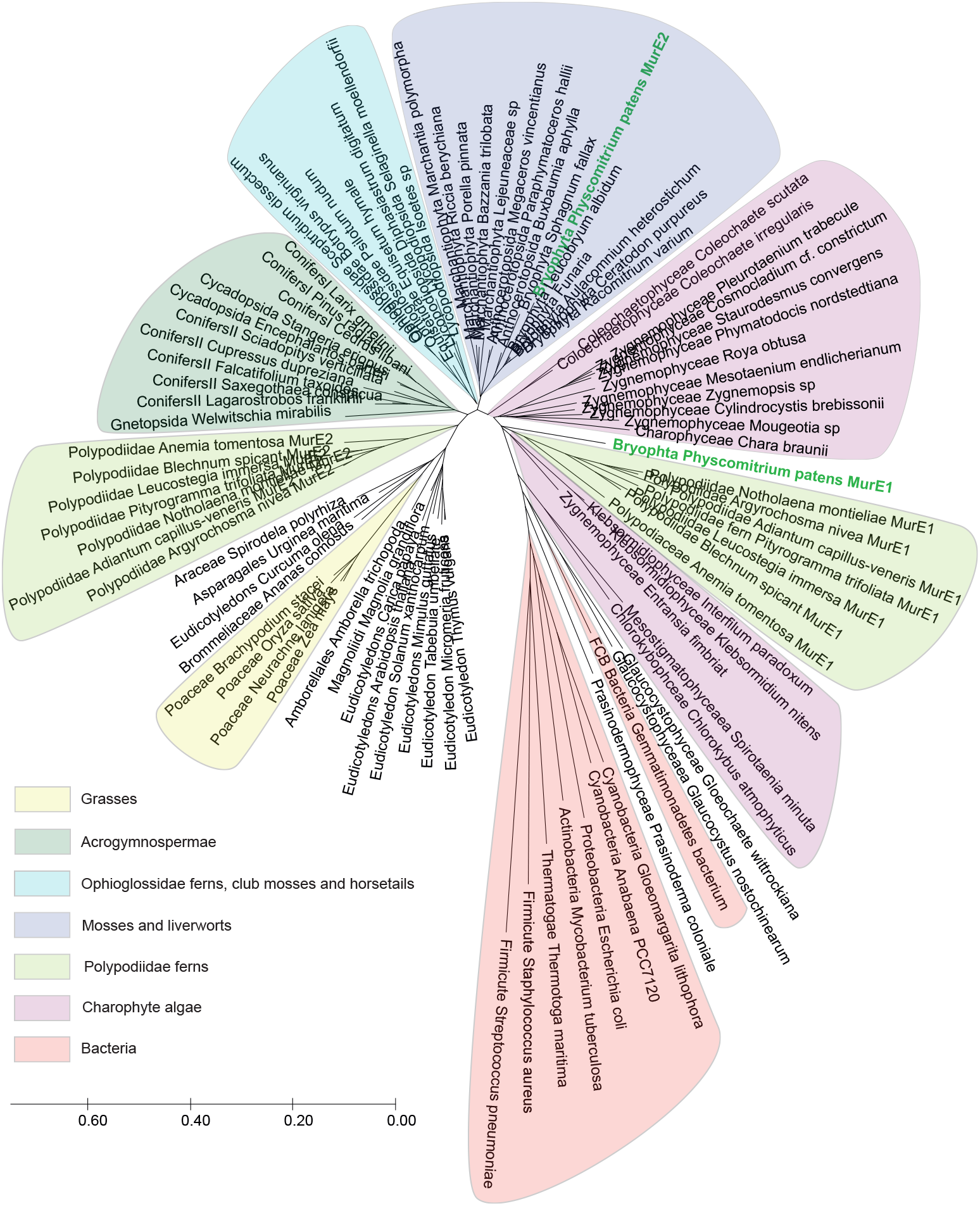
Evolutionary relationship of both PpMurE proteins to selected MurE homologs. *P. patens,* as well as many ferns in the Polypodiidae, encodes two MurE homologs: PpMurE1 and PpMurE2, labelled in green. Different taxonomic groups are boxed to highlight the relationship of the *P. patens* proteins to bacterial, algal and streptophyte phylla. Sequences, except *P. patens*, were sourced from the ONEKP database and selected to represent each group (Leebens-Mack et al 2019, Carpenter et al 2019). The evolutionary history was inferred using the Minimum Evolution method (Rzhetsky and Nei,1992) and computed using MegaX software (Kumar et al 2018). The evolutionary distances, are in the units of the number of amino acid substitutions per site. PpMurE1 and the shorter Polypodiidae fern ‘MurE1’ homologs are evolutionarily closer to charophyte algae than land plants. PpMurE2 is closer to most marchantiophytes and other bryophytes, which lack a second MurE homolog, whereas the longer Polypodiidae ‘MurE2’ are closer to the Acrogymnsopermae (conifers). PpMurE2 primarily differs from PpMurE1 in comprising a long, relatively unstructured extension at the amino terminus and a short carboxy terminal extension. The former is considerably longer than a conventional transit peptide (290 residues longer than typical bacterial homologs, compared to 94 residues for PpMurE1) and is common to most seed plant MurE-like proteins, as well as some streptophyte algal and bryophyte MurE homologs. The extended amino terminus is typically proline-rich in the amino terminal residues, being more glycine-rich in lower orders, and, in the later residues, more conserved within different plant divisions. The carboxy terminal extension (24 residues in PpMurE2 beyond a consensus streptophyte DDREECREAL motif in PpMurE1 (Supplemental Figure S9) is more highly conserved, with a consensus sequence (DDREECREALQXVDXLHXAGIDTFESPWRXPESX) that is common to most streptophyte MurE homologs, although streptophyte algae lack the terminal PESX. However, where there are two distinct MurE homologs, as there are for *P. patens* and some ferns, this carboxy terminal extension is typically absent from the shorter MurE homologs and these proteins appear to have de-evolved to more closely resemble their bacterial counterparts. The retention of a DNPR motif is common not only to the non-seed plants but also most seed plant MurE homologs, with the similarly charged DNPK also being common, and the Poaceae and a few Pinaceae being notable exceptions (DNPA and DNSR, respectively). In contrast to *P. patens* and the Polypodiidae ferns, many in the same and closely related phylla, including the Acrogymnospermae (Lin et al., 2017) do not have two candidate MurE homologs yet they encode most of the peptidoglycan synthesis enzymes.

